# 5′ transgenes drive leaky expression of 3′ transgenes in inducible bicistronic vectors

**DOI:** 10.1101/2022.04.23.489261

**Authors:** Yasuyuki Osanai, Yao Lulu Xing, Kenta Kobayashi, Jihane Homman-Ludiye, Amali Cooray, Jasmine Poh, Nobuhiko Ohno, Tobias D. Merson

**Affiliations:** Australian Regenerative Medicine Institute, 15 Innovation Walk, Monash University, Clayton, Victoria, 3800, Australia.; Department of Anatomy, Division of Histology and Cell Biology, School of Medicine, Jichi Medical University, Shimotsuke, Japan.; Section of Viral Vector Development, National Institute for Physiological Sciences, Okazaki, Japan.; SOKENDAI (The Graduate University for Advanced Studies), Okazaki, Japan.; Division of Ultrastructural Research, National Institute for Physiological Sciences, Okazaki, Japan.

**Keywords:** Gene expression, Bicistronic vector, Leaky expression, FLEX, DIO, lox-STOP-lox, AAV

## Abstract

Molecular cloning techniques enabling contemporaneous expression of two or more protein-coding sequences in a cell type of interest provide an invaluable tool for understanding the molecular regulation of cellular functions. DNA recombination employing the Cre-lox system is commonly used as a molecular switch for inducing the expression of recombinant proteins encoded within a bicistronic cassette. In such an approach, the two protein-coding sequences are separated by a 2A peptide or internal ribosome entry site (IRES), and expression is designed to be strictly Cre-dependent by using a *lox*-STOP-*lox* cassette or flip-excision (FLEX) switch. However, low-level or ‘leaky’ expression of recombinant proteins is often observed in the absence of Cre activity, potentially compromising the utility of this approach. To investigate the mechanism of leaky gene expression, we generated *pCAG-lox-GFP-STOP-lox-Transgene A-2A-Transgene B* vectors, which are designed to express nuclear-targeted GFP in the absence of Cre, and express both transgenes A and B after Cre-mediated recombination. We found that cells transfected with these bicistronic vectors exhibited low-level Cre-independent expression specifically of the transgene positioned 3′ of the 2A peptide. We observed similar results in vivo by viral transduction of the adult mouse cerebral cortex with AAV-mutagenesis of putative transcription factor binding sites that the 5′ transgene confers promoter-like activity that drives expression of the 3′ transgene. Finally, we demonstrate that inclusion of an additional lox-STOP-lox cassette between the 2A sequence and 3′ transgene dramatically reduces the extent of Cre-independent leaky gene expression. Our findings highlight that caution should be applied to the use of Cre-dependent bicistronic constructs when tight regulation of transgene expression is desired and provide a guide to preventing leaky gene expression when the expression of more than one protein is required.

## Introduction

Cell type-specific bicistronic expression systems are powerful tools to investigate cell morphology and physiology (Wang, 2009; Yamamoto et al., 2009). To express recombinant proteins in a cell type-specific manner, a transgene regulated by a cell type-specific promoter or a combination of cell type-specific Cre transgenic mice with *lox-*STOP*-lox* or FLEX vectors is widely used. The *lox*-STOP-*lox* cassette comprises two identical unidirectional *lox* sequences that flank a STOP sequence (poly(A) signal) designed to terminate transcription (Lakso et al., 1992) (Madisen et al., 2010). Inserting a fluorescent reporter gene, such as tdTomato immediately before the STOP sequence and inserting another reporter, such as GFP after the *lox*-STOP-*lox* cassette enables one to distinguish between cells that have exhibited Cre activity (tdTomato^−^/GFP^+^) versus those that have not been exposed to Cre (tdTomato^+^/GFP^−^) (Werdien et al., 2001) (Muzumdar et al., 2007). An alternative strategy for Cre-dependent expression of transgenes is to use a flip-excision (FLEX) switch, also known as a double-floxed inverted open reading frame (DIO). FLEX switches typically use two pairs of non-compatible *lox* sequence variants, such as a pair of *loxP* sites and a pair of either *lox2272*, *loxN* or *lox5171*, to flip the orientation of a protein-coding sequence (CDS) from antisense to sense orientation upon Cre-mediated flip/excision (Lee and Saito, 1998) (Richier and Salecker, 2015). Both *lox-*STOP*-lox* sequences and FLEX switches have been used to drive the overexpression of transgenes in cells that express Cre recombinase in a cell type-specific manner (Kim et al., 2018). The smaller size of the FLEX switch (<300 bp) compared to most *lox-STOP-lox* cassettes (∼1000 bp) make it particularly attractive for use in AAV vectors where cargo size is limited to <5 kb including promoter, CDS and poly(A) sequence (Wu et al., 2010).

In many instances, the expression of multiple transgenes under the control of a single promoter is desired. One approach is to generate a fusion protein by concatenating the protein-coding sequences of two or more proteins into a single CDS. Fusion proteins are useful for identifying the subcellular localization of a target protein, for example by expressing a protein of interest with an expression tag such as GFP. However, this strategy cannot be used if the expression of proteins with different subcellular localizations is required or if the generation of the fusion protein alters the function of either protein. In this case, the CDS encoding two or more proteins can be separated by an internal ribosomal entry site (IRES) or self-cleaving 2A peptide. Since the IRES sequence is longer than the 2A peptide sequence (∼500 bp vs 66 bp), and the expression level of the 3′ CDS is typically significantly lower than that of the 5′ CDS expression in IRES constructs (Mizuguchi et al., 2000), the use of 2A peptides has become the preferred method to express multiple genes under the control of a single promoter.

For maximal utility, an inducible expression system should allow for stringent control of transgene expression. Yet leaky expression of transgenes in the absence of an induction signal is frequently observed (Fischer et al., 2019; Kallunki et al., 2019; Lavin et al., 2020). For inducible transgene expression systems that employ Cre recombinase as the induction signal, low levels of Cre-independent expression may be tolerable, depending upon the proteins being expressed. Low-level GFP expression, for instance,may be acceptable in an animal injected with an adeno-associated virus designed to express the light-sensitive neuronal silencer archaerhodopsin and GFP (AAV-*FLEX-ArchT-GFP*). However, in some cases, even very low levels of leaky transgene expression can have a significant impact on cell function and on the interpretation of experimental data, as illustrated by the following three examples. First, leaky expression of diphtheria toxin fragment A (DTA), a transgene widely used as a suicide gene to ablate cells of interest (Lee et al., 1998; Stanger et al., 2007), would be unacceptable since a single molecule of DTA is sufficient to kill a cell (Yamaizumi et al., 1978). Second, the loss of astrocyte-specific expression of mCherry in AAV-*GFAP-NeuroD1-T2A-mCherry* transfected cells, as observed by Wang and colleagues, could lead to the incorrect conclusion that glial cells can transdifferentiate to neurons (Wang et al., 2021). Third, attempts to identify synaptically connected cells by infecting cells with an EnvA-pseudotyped recombinant rabies virus can be hampered by leaky expression of TVA in Cre-negative cells due to the strength of the EnvA-TVA interaction (Lavin et al., 2020) (Federspiel et al., 1994) (Seidler et al., 2008; Wall et al., 2010).

Understanding the cause of leaky transgene expression is key to improving the stringency of inducible expression systems. It has been demonstrated that Cre-independent expression of transgenes encoded by bicistronic FLEX vectors can be reduced by repositioning the ATG start codon to a region 5′ of the *loxP* site just downstream of the promoter (Wall et al., 2010) (Fischer et al., 2019). However, approaches to eliminate Cre-independent leaky expression are yet to be fully explored (Lavin et al., 2020; Wang et al., 2021).

In this study, we demonstrate that transgenes positioned 3′ of a 2A sequence in Cre-inducible bicistronic expression constructs are prone to Cre-independent expression. The leaky expression is not derived from a typical promoter, but arises from promoter-like activity of the transgene that is positioned 5′ of the 2A sequence. We demonstrate that introducing a second *lox*-STOP-*lox* sequence, using non-compatible *lox* sites, between the 2A sequence and 3′ CDS, eliminates leaky expression of the 3′ transgene. Our findings have significant implications for transgenesis and gene therapy studies where the optimal design of constructs to precisely control the expression of multiple protein-coding sequences under the regulatory control of a single promoter is desired.

## Results

### Leaky expression of the 3′ transgene in Cre-inducible bicistronic expression vectors

To investigate factors that influence Cre-independent gene expression of Cre regulated gene expression vectors, we generated a *lox*-STOP-*lox* vector (*pCAG-loxP-nGFP-STOP-loxP-rtTA-P2A-memCherry*, hereafter denoted *CAG-LGL-rtTA-mC*) designed to express nuclear-targeted GFP (nGFP) without Cre activity and both the reverse tetracycline transactivator (rtTA) and membrane-targeted mCherry (mC) following Cre activity (**Figure 1A**). Co-expression of rtTA and mC following Cre activity was enabled by creating a bicistronic expression cassette wherein each CDS was separated by a P2A peptide. The construct was transfected into HEK 293T cells either in the presence or absence of *pCAG-Cre*. The cells correctly expressed nGFP in the Cre (-) condition and expressed mCherry in the Cre (+) condition (**Figure 1B**). However, approximately 7 % of the cells transfected with *CAG-LGL-rtTA-mC* expressed mC even in Cre (-) condition (**Figure 1C, 1D**). The leakage of mC (8.2%) was further confirmed by flow cytometry (**Figure 1E**).

**Figure 1.**
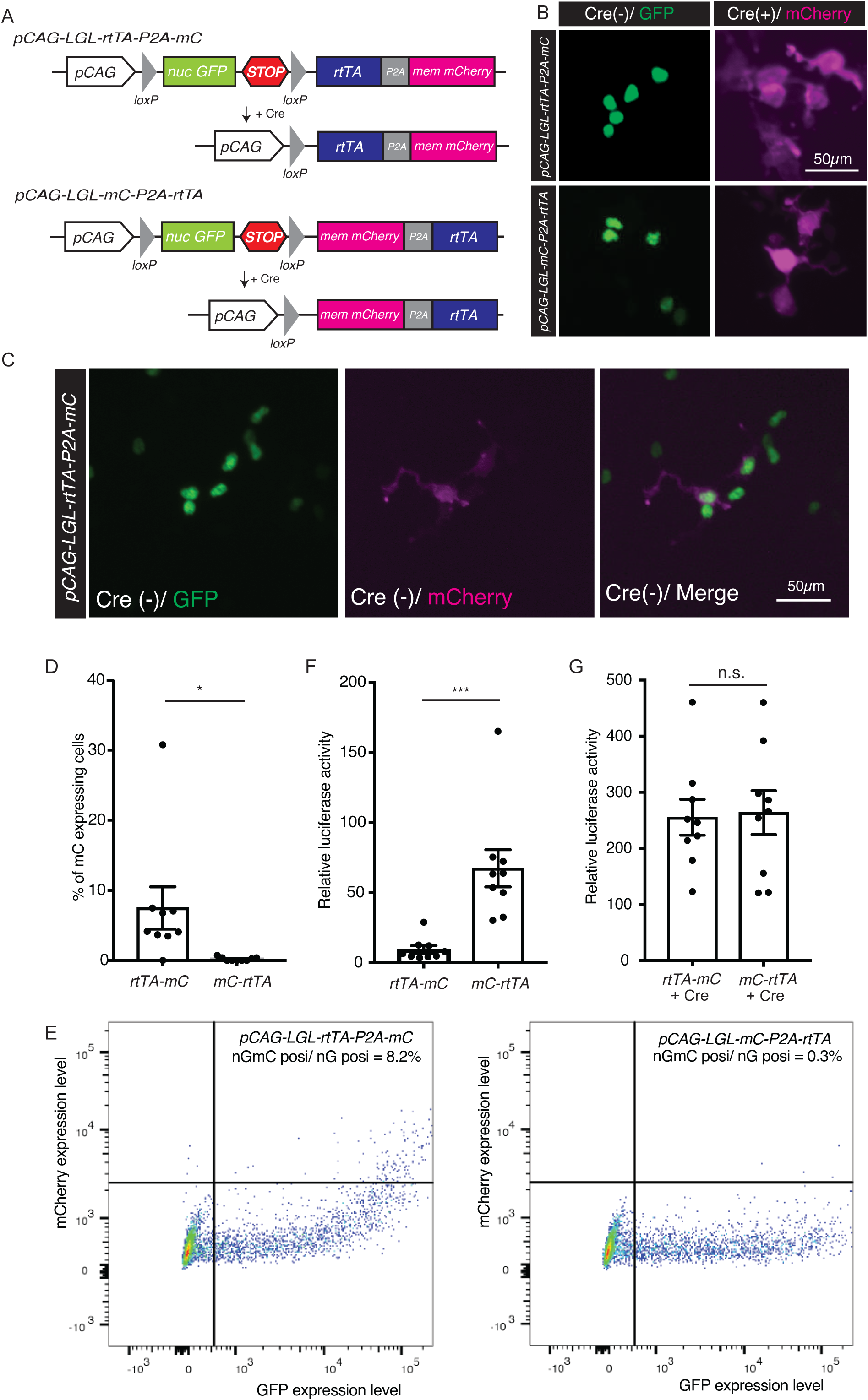
Transgenes located in the 3′ region of bicistronic vectors are leaky. (A) Scheme of *CAG-LGL-rtTA-mC* or *CAG-LGL-mC-rtTA* vector. The difference between these two vectors is the position of rtTA and mC. (B) HEK293T cells transfected with *CAG-LGL-rtTA-mC* (top) or *CAG-LGL-mC-rtTA* (bottom) expressing nucleus-targeted GFP in Cre (-) condition or membrane-targeted mCherry in Cre (+) condition. Scale bar, 50 µm. (C) *CAG-LGL-rtTA-mC*-transfected HEK293T cells expressing leaky mC in Cre (-) condition. Scale bar, 50 µm. (D) Comparison of the percentage of transfected cells expressing leaky mC between *CAG-LGL-rtTA-mC*- and *CAG-LGL-mC-rtTA*-transfected cells in Cre (-) condition (*rtTA-mC* 7.49 ± 3.01 %, *mC-rtTA* 0.19 ± 0.09 %, n = 9 wells for three independent experiment). (E) Flow cytometric analysis of HEK cells transfected with *CAG-LGL-rtTA-mC* (left) or *CAG-LGL-mC-rtTA* (right); 8.2% or 0.3% of GFP-positive *CAG-LGL-rtTA-mC or CAG-LGL-mC-rtTA-* transfected cells expressed leaky mCherry, respectively. (F) Comparison of relative luciferase activity between cells transfected with *CAG-LGL-rtTA-mC* or *CAG-LGL-mC-rtTA* in Cre (-) condition and those co-transfected with *pTRE-Luciferase* (*rtTA-mC* 9.43 ± 2.67, *mC-rtTA*67.3 ± 13.3, n = 9 wells from three independent experiment). (G) Relative luciferase activity of *CAG-LGL-rtTA-mC*- or *CAG-LGL-mC-rtTA*-transfected cells and cells co-transfected with *pTRE-Luciferase* and *pCAG-Cre* (*rtTA-mC* 255.6 ± 10.6, *mC-rtTA* 263.7 ± 13.0, n = 9 wells from three independent experiment). n.s. p > 0.05, *p < 0.05, ***p < 0.001 using two-tailed Student’s t-test. Data shown as mean ± SEM.

We hypothesized that mC was leaky because the protein coding sequence (CDS) of mC was located far from the STOP cassette. Thus, the order of rtTA and mC was swapped, and *pCAG-loxP-nG-STOP-loxP-mC-P2A-rtTA* (hereafter referred as *CAG-LGL-mC-rtTA*) was generated (**Figure 1A**). The number of cells expressing leaky mC significantly decreased in *CAG-LGL-mC-rtTA*-transfected cells as compared to that in *CAG-LGL-rtTA-mC*-transfected cells (*rtTA-mC* 7.49 ± 3.01 %, *mC-rtTA* 0.19 ± 0.09 %, p = 0.03, n = 9, **Figure 1D**). The reduced leaky mC expression in *CAG-LGL-mC-rtTA*-transfected cells was further confirmed via flow cytometry (0.3%, **Figure 1E**). Next, we analysed rtTA expression in *CAG-LGL-rtTA-mC* and *CAG-LGL-mC-rtTA* via luciferase assay using *pTRE-Luciferase* plasmid, which expresses luciferase in response to rtTA expression (see *Materials and Methods*). In contrast to mC leakage, rtTA expression of *CAG-LGL-rtTA-mC*-transfected cells was finely regulated while *CAG-LGL-mC-rtTA*-transfected cells expressed leaky rtTA without Cre activity (*rtTA-mC* 9.43 ± 2.67, *mC-rtTA* 67.3 ± 13.3, p = 0.0006, n = 9, **Figure 1F**). As shown in Figure 1D-1F, the leakage expression pattern was reversed by swapping the positions of rtTA and mC in Cre (-) condition, indicating that the transgene located at 3′ position was leaky. Both constructs correctly expressed mC and rtTA on *pCAG-Cre* transfection (**Figure 1B**) (Relative luciferase activity; *rtTA-mC*255.6 ± 10.6, *mC-rtTA* 263.7 ± 13.0, p = 0.86, n = 9, **Figure 1G**). As the leaky expression levels of mC and rtTA fused via P2A were quite different within the transfected cells, the expression of rtTA and mC in Cre (-) condition seemed to be “leaky” as compared to the result of Cre-independent recombination. Thus, the transgene located at the 3′ region of bicistronic vector was leaky in Cre-inducible vector.

### Changing the promoter does not prevent leaky expression

We checked whether swapping the promoter prevents the leaky expression of the 3′ CDS. The CAG promoter was replaced with myelin basic protein (MBP) promoter, which is active specifically in oligodendrocytes, a type of glial cell in the central nervous system of vertebrates. *MBP-LGL-rtTA-mC* and *MBP-LGL-mC-rtTA* were transfected into the oligodendroglial cell line, CG4 (Louis et al., 1992). The leaky expression of mC was lower in *MBP-LGL-mC-rtTA*-transfected cells as compared to that in *MBP-LGL-rtTA-mC*-transfected cells without Cre activity (*rtTA-mC* 2.79 ± 0.78 %, *mC-rtTA* 0.14 ± 0.14 %, p = 0.003, n = 12, **Figure 2A**). Leaky mCherry expression in *MBP-LGL-rtTA-mC*-transfected cells was further confirmed via flow cytometry (*rtTA-mC* 3.3%, *mC-rtTA* 0%, **Figure 2B**). Predictably, the leaky expression level of rtTA was higher in *MBP-LGL-mC-rtTA-*transfected cells as compared with *MBP-LGL-rtTA-mC* transfected cells *(rtTA-mC* 1.19 ± 0.05, *mC-rtTA* 6.71 ± 1.29, p = 0.00007, n = 33, **Figure 2C**) while the expression level of rtTA was similar between *MBP-LGL-rtTA-mC* and *MBP-LGL-mC-rtTA* on transfection with *pCAG-Cre* (*rtTA-mC* 164.3 ± 28.7, *mC-2A-rtTA* 129.2 ± 21.4, p = 0.33, n = 24, **Figure 2D**).

**Figure 2.**
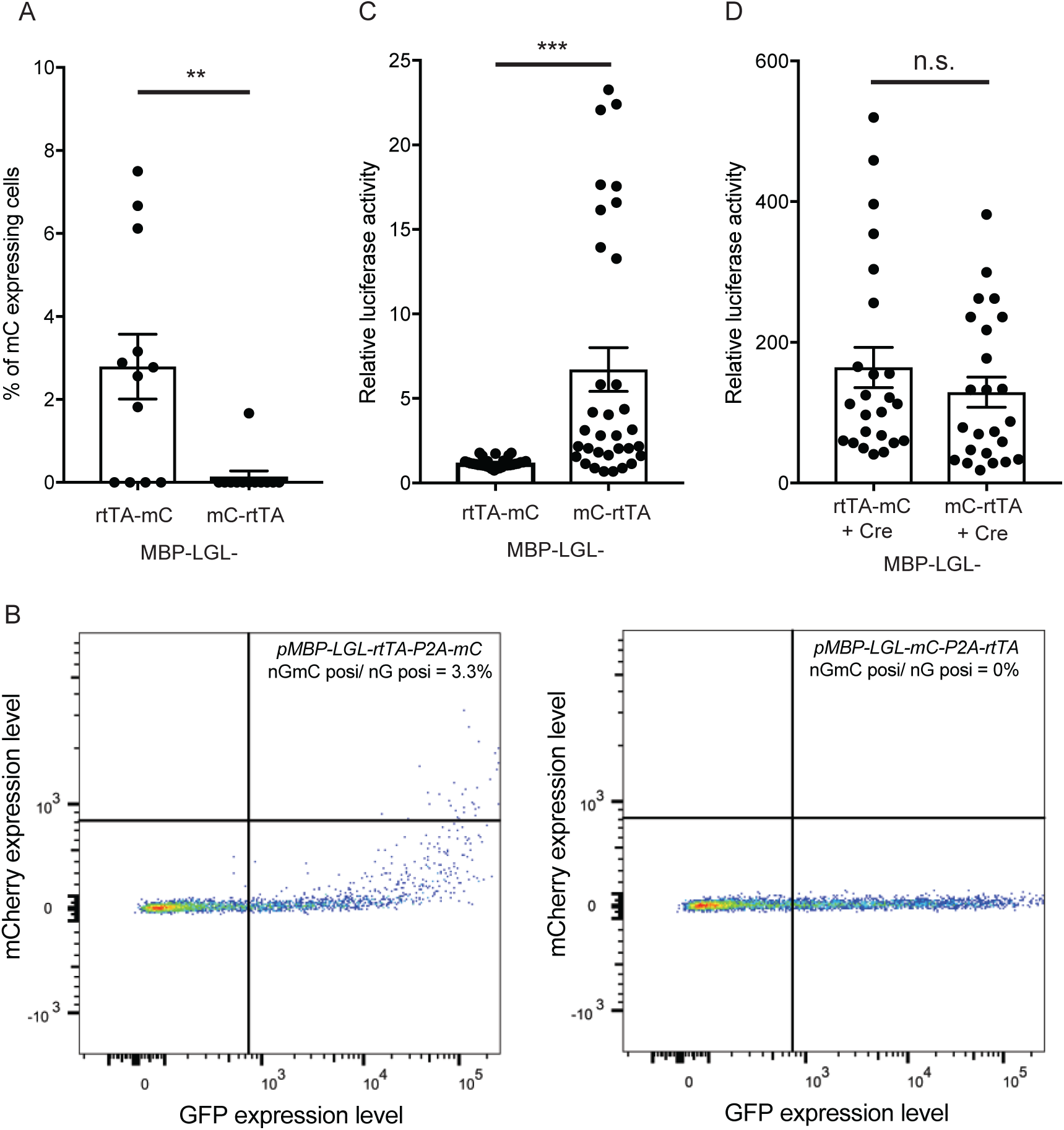
Inducible bicistronic vector with cell type-specific promoter expresses leaky 3′ transgene. (A) Comparison of the percentage of transfected cells expressing leaky mC between *MBP-LGL-rtTA-mC*- or *MBP-LGL-mC-rtTA*-transfected cells in Cre (-) condition (*rtTA-mC* 2.79 ± 0.78 %, *mC-rtTA* 0.14 ± 0.14 %, n = 12 wells from four independent experiments). (B) Flow cytometric analysis of CG4 cells transfected with *MBP-LGL-rtTA-mC* or *MBP-LGL-mC-rtTA*; 3.3% or 0% of GFP (+) cells of *MBP LGL-rtTA-mC-* or *MBP-LGL-mC-rtTA*-transfected cells expressed leaky mCherry, respectively. (C) Comparison of the relative luciferase activity between cells transfected with *MBP-LGL-rtTA-mC* or *MBP-LGL-mC-rtTA* in Cre (-) condition and those co-transfected with *pTRE-Luciferase*. (*rtTA-mC* 1.19 ± 0.05, *mC-rtTA* 6.71 ± 1.29, n = 33 wells from 11 independent experiments). (D) Relative luciferase activity of *MBP-LGL-rtTA-mC* transfected cells or *MBP-LGL-mC-rtTA* transfected cells, co-transfected with *pCAG-Cre* and *pTRE-Luciferase* (*rtTA-mC* 164.3 ± 28.7, *mC-rtTA* 129.19 ± 21.4, n = 24 wells from four independent experiments). ns. p > 0.05, **p < 0.01, ***p < 0.001 using two-tailed Student’s t-test. Data shown as mean ± SEM.

To check whether another set of transgenes in a bicistronic inducible vector expresses 3′ transgene leakage, we generated *MBP-LGL-diphtheria toxin receptor (DTR)-IRES-tdTomato* that expresses DTR and tdTomato when transfected into the cells with *pCAG-Cre*. We found that *MBP-LGL-DTR-IRES-tdTomato-*transfected cells expressed leaky tdTomato without Cre-activity (8.9%, **Figure S1**). In contrast, *MBP-LGL-mC-P2A-DTR-transfected* cells did not express leaky mC without Cre activity (**Figure S1**). Taken together, the leakage of 3′ transgene seemed to be driven by 5′ CDS and not by general promoters.

### Leaky expression in cells transfected with promoter-removed constructs

We hypothesized that 5′ transgene CDSs drive the expression of 3′ transgene. Transgenes located after rtTA CDS or mCherry CDS can be expressed without general promoters. To check whether the leaky protein expression can be observed without general promoters, MBP promoter and STOP cassette were removed from the plasmids by enzymatic digestion (**Figure 3A**), and this was followed by gel extraction to obtain *rtTA-2A-mC* and *mC-2A-rtTA* segments. These *rtTA-2A-mC* and *mC-2A-rtTA* segments were transfected into CG4 cells, and mC or rtTA expression was analysed. Interestingly, some *rtTA-2A-mC*-transfected cells expressed mC without the promoter (**Figure 3B**), and the number of mCherry-expressing cells in *rtTA-2A-mC* transfected cells was significantly higher than that in mC-2A-rtTA transfected cells (rtTA-P2A-mC 4.15 ± 1.07 cells/well, *mC-P2A-rtTA* 0.25 ± 0.11 cells/well, 199 p = 0.0009, n = 20, **Figure 3C**). Notably mCherry-expressing cells did not co-express nGFP (**Figure 3B**), confirming no general promoters in this segment. As expected, rtTA expression was lower in *rtTA-2A-mC*-transfected cells as compared to that in *mC-2A-rtTA*-transfected cells (*rtTA-2A-mC* 2.55 ± 0.35, *mC-2A-rtTA* 7.18 ± 1.84, p = 0.02, n = 18, **Figure 3D**). These data indicated that the CDS located in the 3′ region of bicistronic constructs was expressed without a promoter. These results suggest that constructs with multiple transgenes can be leaky, even if the cell-specific promoter or Cre is inactive in transfected cells.

**Figure 3.**
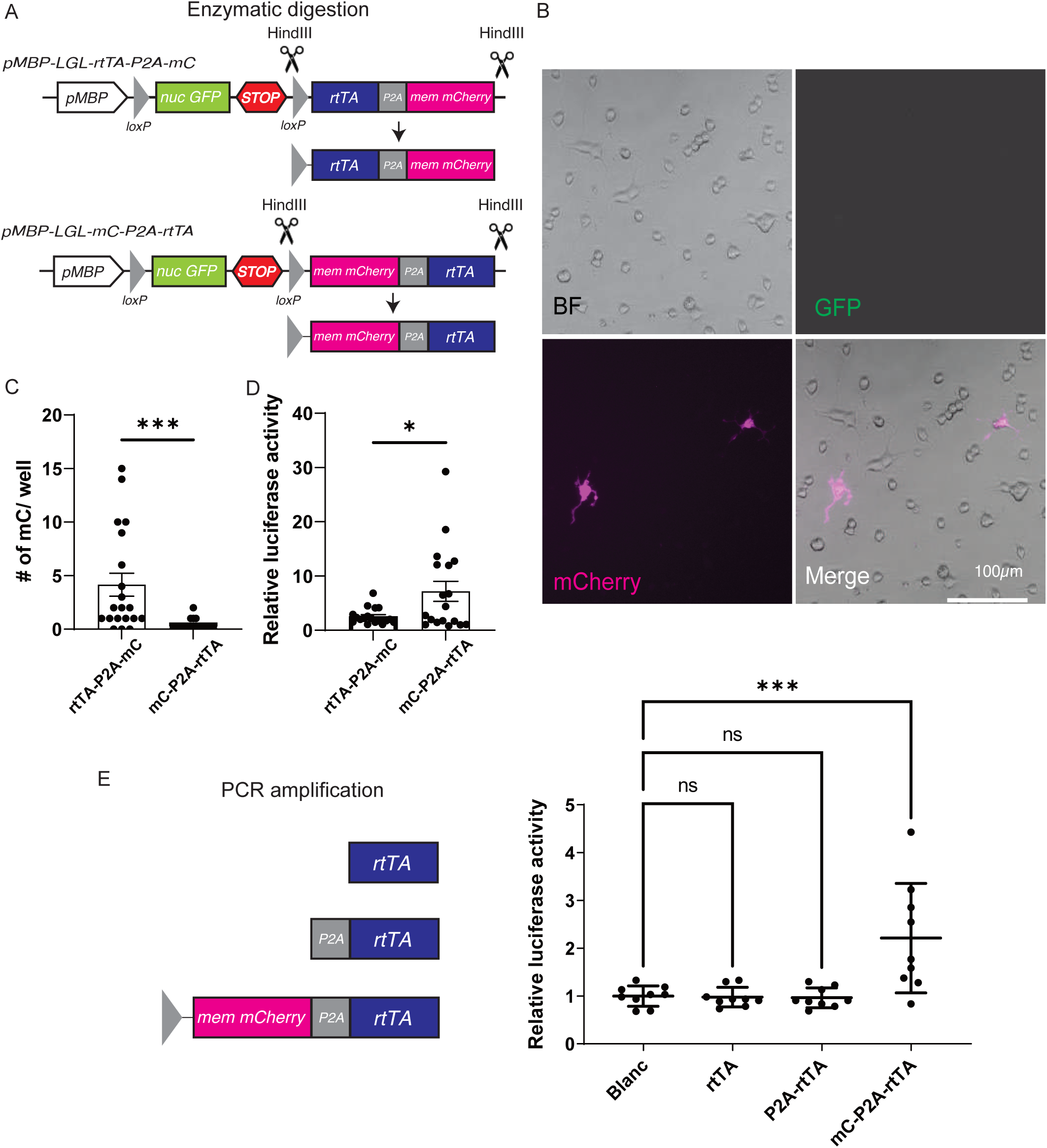
Leaky expression was observed without a general promoter. (A) Schematic diagram of the preparation of *rtTA-P2A-mC* and *mC-P2A-rtTA* segments. (B) *rtTA-P2A-mC* segment-transfected cells expressing mCherry. (C) Comparison of the percentage of transfected cells expressing leaky mC between *rtTA-P2A-mC* segment- or *mC-P2A-rtTA* segment-transfected cells in Cre (-) condition. (*rtTA-P2A-mC* 4.15 ± 1.07 cells/well, *mC-P2A-rtTA* 0.25 ± 0.11 cells/well, n = 20 wells from six independent experiment). (D) Comparison of the relative luciferase activity between cells transfected with *rtTA-P2A-mC* or *mC-P2A-rtTA* segment in Cre (-) condition, co-transfected with *pTRE-Luciferase* (*rtTA-2A-mC* 2.55 ± 0.35, *mC-2A-rtTA* 7.18 ± 1.84, n = 18 wells from six independent experiments). (E) Left figure; Schematic diagram of PCR-amplified rtTA segment, *P2A-rtTA* segment, and *mC-P2A-rtTA* segment. Right figure depicts the relative luciferase activity of cells transfected with PCR-amplified *rtTA* segment, *P2A-rtTA* segment, or *mC-P2A-rtTA* segment and those co-transfected with *pTRE-Luciferase* (Blanc 1.00 ± 0.07, *rtTA* segment 0.98 ± 0.07, *P2A-rtTA* segment 0.96 ± 0.07, *mC-P2A-rtTA segment* 2.21 ± 0.38, p = 0.001, n =9 wells from three independent experiments). n.s. p > 0.05, *p < 0.05, ***p < 0.001, two-tailed Student’s t-test for Figures 3C and 3D or one-way ANOVA with Dunnett’s test for Figure 3E. Data shown as mean ± SEM.

To investigate the possibility that P2A sequence is responsible for the leaky expression, PCR-amplified *rtTA* segment, *P2A-rtTA* segment, and *mC-P2A-rtTA* segment were transfected into CG4 cells and the rtTA expression level was compared. Cells transfected with either the *rtTA* segment or *P2A-rtTA* segment did not express rtTA whereas cells transfected with the *mC-P2A-rtTA* segment expressed high levels of rtTA (Blank 1.00 ± 0.07, *rtTA* segment 0.98 ± 0.07, *P2A-rtTA* segment 0.96 ± 0.07, *mC-P2A-rtTA* segment 2.21 ± 0.38, p = 0.001, **Figure 3E**). Taken together, the leaky expression of 3′ region CDS in bicistronic constructs was not driven by P2A but by the 5′ region CDS.

### Multiple transcription binding sites are present in both mC and rtTA sequences

Assembly of a transcription initiation complex comprising transcription factors and RNA polymerase is a pre-requisite for gene expression. We checked whether both mCherry CDS and rtTA CDS had transcription factor (TF)-binding sites. Using JASPAR TF motif, we found multiple TF binding regions in both mC CDS and rtTA CDS (**Figure S2A,S2B**). To assess whether the removal of TF binding sites changes the leaky expression, we performed silent mutation of 20 individual codons in the mCherry CDS to mutate putative 20 TF binding motifs without altering the amino acid sequence (**Figure S2A, S2B**). We selected those TFs that have been identified as transcriptional activators and which have also been shown to be expressed in oligodendrocyte progenitor cells, origin of the CG4 cell line. The TF-modified mCherry CDS was used in place of the parent mCherry CDS in order to generate a new construct denoted *MBP-LGL-TF modified-mC-P2A-rtTA* (here after *MBP-LGL-TFmC-rtTA*). We observed that *MBP-LGL-TFmC-rtTA* correctly expressed GFP prior to Cre recombination, and it correctly expressed mC following Cre recombination in CG4 cells (**Figure S2C**). Unexpectedly, we found that the leaky expression level of rtTA in *MBP-LGL-TFmC-rtTA*-transfected cells was higher than that in normal *MBP-LGL-mC-rtTA*-transfected cells (*mC-rtTA* 15.7 ± 1.57, *TFmC-rtTA* 25.0 ± 3.73, p = 0.03, n = 12, **Figure S2D**), even though the expression level of rtTA was similar after Cre recombination (*mC-rtTA* 129.9 ± 35.1, *TFmC-rtTA* 102.4 ± 22.1, p = 0.52, n = 6, **Figure S2D**). One possible explanation could be that the modification of TF binding sites generated other TF binding sites, leading to strong promoter activity of mC. We also generated another version of *MBP-LGL-TFmC-rtTA*, which removed other sets of TF binding sites. However, the construct also showed higher leaky rtTA expression as compared to normal *MBP-LGL-mC-rtTA* (data not shown). These results suggest that silent mutation of codons in the 5′ region CDS alters the degree of leaky expression of transgene encoded by the 3′ CDS.

### Inserting transgenes into the genome reduces leaky transgene expression

Although our results indicated that transient transfection of inducible bicistronic vector led to a significant amount of leakage, some studies on transgenic mice with bicistronic cassette indicated that the leakage is not observed often (Madisen et al., 2012) (Sciolino et al., 2016). We reasoned that transgenes integrated into the genome would not express a high amount of leakage. To analyse whether integration of the bicistronic construct reduces leaky transgene expression, we generated stable cell lines HEK-*CAG-LGL-rtTA-P2A-mC* (hereafter, HEK*-rtTA-mC*) and HEK-*CAG-LGL-mC-P2A-rtTA* (hereafter, HEK*-mC-rtTA*). Leaky 3′ transgene expression showed a similar trend with no statistical significance (For mC leakage; HEK-*rtTA-mC* 0.30 ± 0.14 %, HEK-*mC-rtTA* 0.05 ± 0.03 %, p = 0.10, n = 9, **Figure 4A, 4B**; for rtTA, HEK-*rtTA-mC* 1.67 ± 0.30, HEK *mC-rtTA* 3.11 ± 0.78, p = 0.11, n = 9, **Figure 4C**). Importantly, mC and rtTA leakage levels were approximately > 20 times lower as compared to those of transient transfection into HEK cells (For mC leakage, transient *CAG-LGL-rtTA-mC* transfection 7.49 ± 3.01 %, HEK-*rtTA-mC* 0.30 ± 0.14 %, p = 0.03, n = 9, **Figure 4D**; for rtTA leakage, transient *CAG-LGL-mC-rtTA* transfection 67.3 ± 13.3, HEK-*mC-rtTA* 3.11 ± 0.78, p < 0.001, n = 9, **Figure 4E**). In the presence of Cre, the stable cell line expressed mC correctly (**Figure 4F**). Stable expression cells expressed a lower amount of rtTA in Cre (+) condition than transiently transfected cells (transient *CAG-LGL*-*rtTA-mC* 255.6 ± 31.9, transient *CAG-LGL-mC-rtTA* 263.7 ± 39.1, HEK-*rtTA-mC* 161.5 ± 45.3, HEK-*mC-rtTA* 125.8 ± 48.5, p = 0.06, n = 9, **Figure 4G**). Although rtTA expression was lower in Cre (+) condition in the stable cell lines, the reduction of rtTA leakage (95.4% reduction, transient *CAG-LGL-mC-rtTA* transfection 67.3 ± 13.3, HEK-*mC-rtTA* 3.11 ± 0.78, **Figure 4E**) was much more significant than that in Cre (+) condition (52.3% reduction, transient *CAG-LGL-mC-rtTA* transfection 263.7 ± 39.1, HEK-*mC-rtTA* 125.8 ± 48.5, **Figure 4G**). These data suggested that the 3′ transgene leakage was more pronounced in transient expression experiments than that in stable expression experiments.

**Figure 4.**
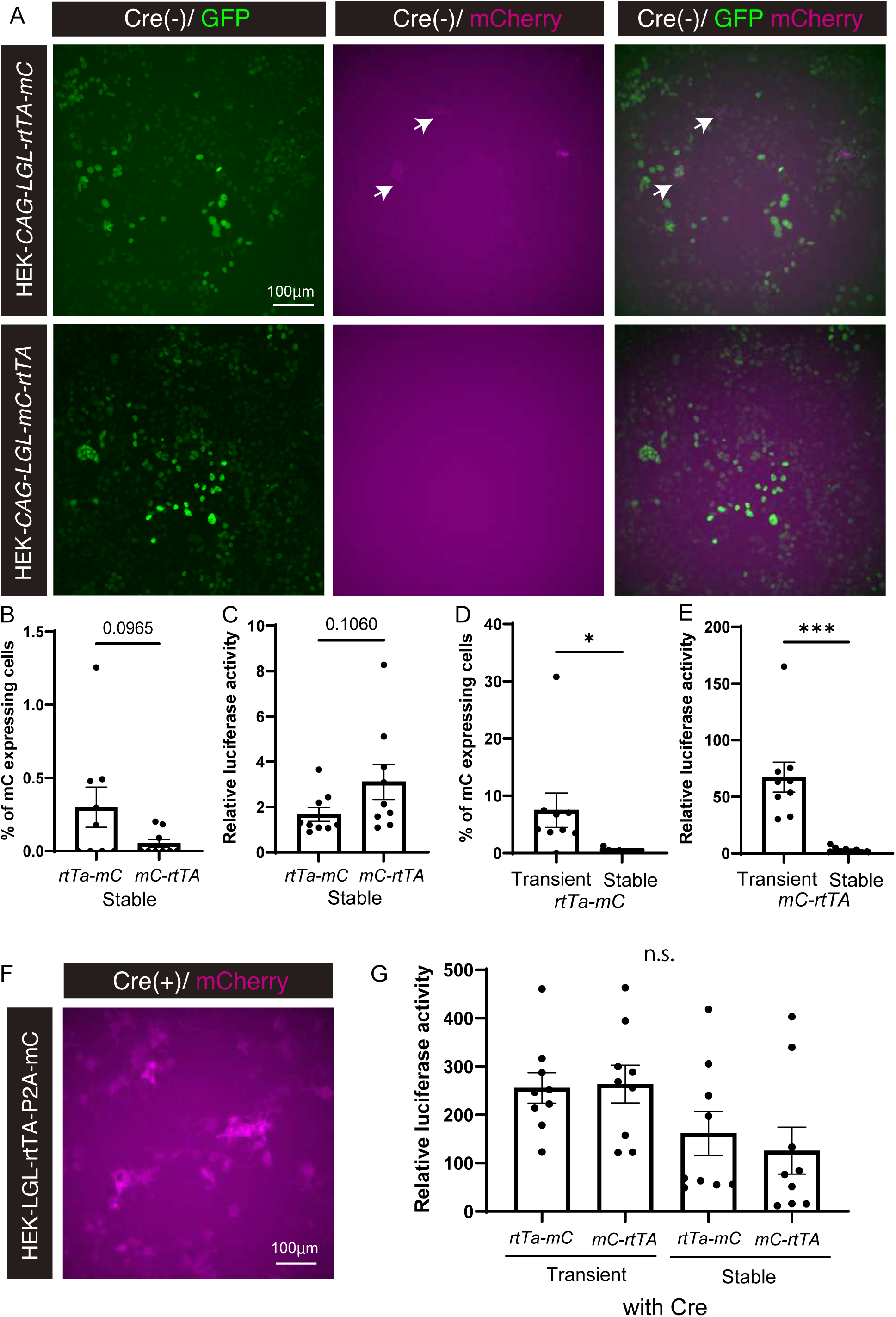
Significant reduction in 3′ transgene leakage in stable cell lines with inducible bicistronic expression. (A) Representative images of HEK-*CAG-LGL-rtTA-mC* cell line (top) or HEK*-CAG-LGL-mC-rtTA* cell line (bottom). Both cell lines correctly expressed nGFP in Cre (-) condition; however, HEK-*CAG-LGL-rtTA-mC* cell line expressed leaky mC (arrows). Scale bar, 100 µm. (B) Comparison of the percentage of cells expressing leaky mC between HEK-*CAG-LGL-rtTA-mC* and HEK-*CAG-LGL-mC-rtTA* cell lines in Cre (-) condition. (*rtTA-mC* 0.30 ± 0.14 %, *mC-rtTA* 0.05 ± 0.03 %, p = 0.0965, n = 9 wells from three independent experiments). (C) Comparison of the relative luciferase activity between HEK-*CAG-LGL-rtTA-mC* and HEK-*CAG*-*LGL-mC-rtTA* cell lines transfected with *pTRE-Luciferase* (*rtTA-mC* 1.67 ± 0.30, *mC-rtTA* 3.11 ± 0.78, p = 0.106, n = 9 wells from three independent experiments). (D) Comparison of the percentage of cells expressing leaky mC between *CAG-LGL-rtTA-mC*-transfected HEK cells and HEK-*CAG-LGL-rtTA-mC* cell line in Cre (-) condition; data sets were obtained from Figure 1D or Figure 4B, respectively (transient *rtTA-mC* 7.49 ± 3.01 %, stable *rtTA-mC* 0.30 ± 0.14 %, n = 9 wells from three independent experiments). (E) Comparison of the relative luciferase activity between *CAG-LGL-mC-rtTA*-transfected HEK cells and HEK-*CAG-LGL-mC-rtTA* cell line in Cre (-) condition those transfected with *pTRE-Luciferase*; data sets were obtained from Figure 1F or Figure 4C, respectively (transient *mC-rtTA* 67.3 ± 13.3, stable *mC-rtTA* 3.11 ± 0.78, n = 9 wells from 3 independent experiments). (F) Photomicrograph of HEK*-CAG-LGL-rtTA-mC* cells transfected with *pCAG-Cre*, expressing mCherry. Scale bar 100µm. (G) Comparison of relative luciferase activity between *CAG-LGL-rtTA-mC* transfected cells, *CAG-LGL-mC-rtTA* transfected cells, HEK-*CAG-LGL-rtTA-mC* cell line and HEK-*CAG-LGL-mC-rtTA* cell line, cells were co-transfected with *pCAG-Cre* and *pTRE-Luciferase* (transient *rtTA-2A-mC* 255.6 ± 31.9, transient *mC-2A-rtTA* 263.7 ± 39.1, stable *rtTA-2A-mC* 161.5 ± 45.3, *mC-2A-rtTA* 125.8 ± 48.5, n = 9 wells from 3 independent experiments), data sets for transient expression were obtained from Figure1G. *p< 0.05, ***p< 0.001 or actual p values are indicated, two-tailed Student’s t-test for Figures 4B-4E, or one-way ANOVA with Tukey test for Figure 4G. Data shown as mean ± SEM.

### STOP cassette insertion into the 3′ region of P2A prevents the leaky expression

We reasoned that inserting another lox-STOP-lox cassette between P2A and 3′ transgene might reduce the leakage. We inserted another lox-STOP-lox site after P2A sequence and generated *MBP-lox2272-nG-STOP-lox2272-mC-P2A-loxP-STOP-loxP-rtTA* (here after *MBP-LGL-mC-LSL-rtTA*) (**Figure 5A**). This construct expressed nG in Cre (-) condition, and it expressed both mC and rtTA in Cre (+) condition (**Figure 5A, 5B**) because both lox2272-nG-STOP-lox2272 and loxP-STOP-loxP sites were independently removed via Cre recombination. Further, we inserted an M3 promoter domain into the MBP promoter, which is known to increase the MBP promoter activity and specificity (Dionne et al., 2016) (Dib et al., 2011) (Farhadi et al., 2003). Cells transfected with the construct did not express leaky mC (no cells expressed mC in 2,130 transfected cells; n = 9 wells). As expected, the leaky rtTA expression level of *MBP-LGL-mC-LSL-rtTA* was significantly lower than that of *MBP-mC-rtTA*-transfected cells in Cre (-) condition (*rtTA-mC* 1.16 ± 0.08, *mC-rtTA* 11.9 ± 1.95, *mC-LSL-rtTA* 1.42 ± 0.12, n = 33, 33, 18; p < 0.001, **Figure 5C**). rtTA expression of *MBP-mC-LSL-rtTA* transfected cells in Cre (+) condition was approximately 30% of normal constructs (*rtTA-mC* 164.3 ± 28.7, *mC-rtTA* 129.2 ± 21.4, *mC-LSL-rtTA* 47.1 ± 8.03, n = 24, 24, 18; p = 0.0033, **Figure 5D**). Although the rtTA expression level was low in Cre (+) condition, the construct is suitable for a transgene that requires both precise Cre-mediated expression and low gene expression, e.g., TVA. These results supported our findings that the 5′ CDS drives 3′ transgene expression, suggesting that the expression could be prevented via additional lox-STOP-lox insertion.

**Figure 5.**
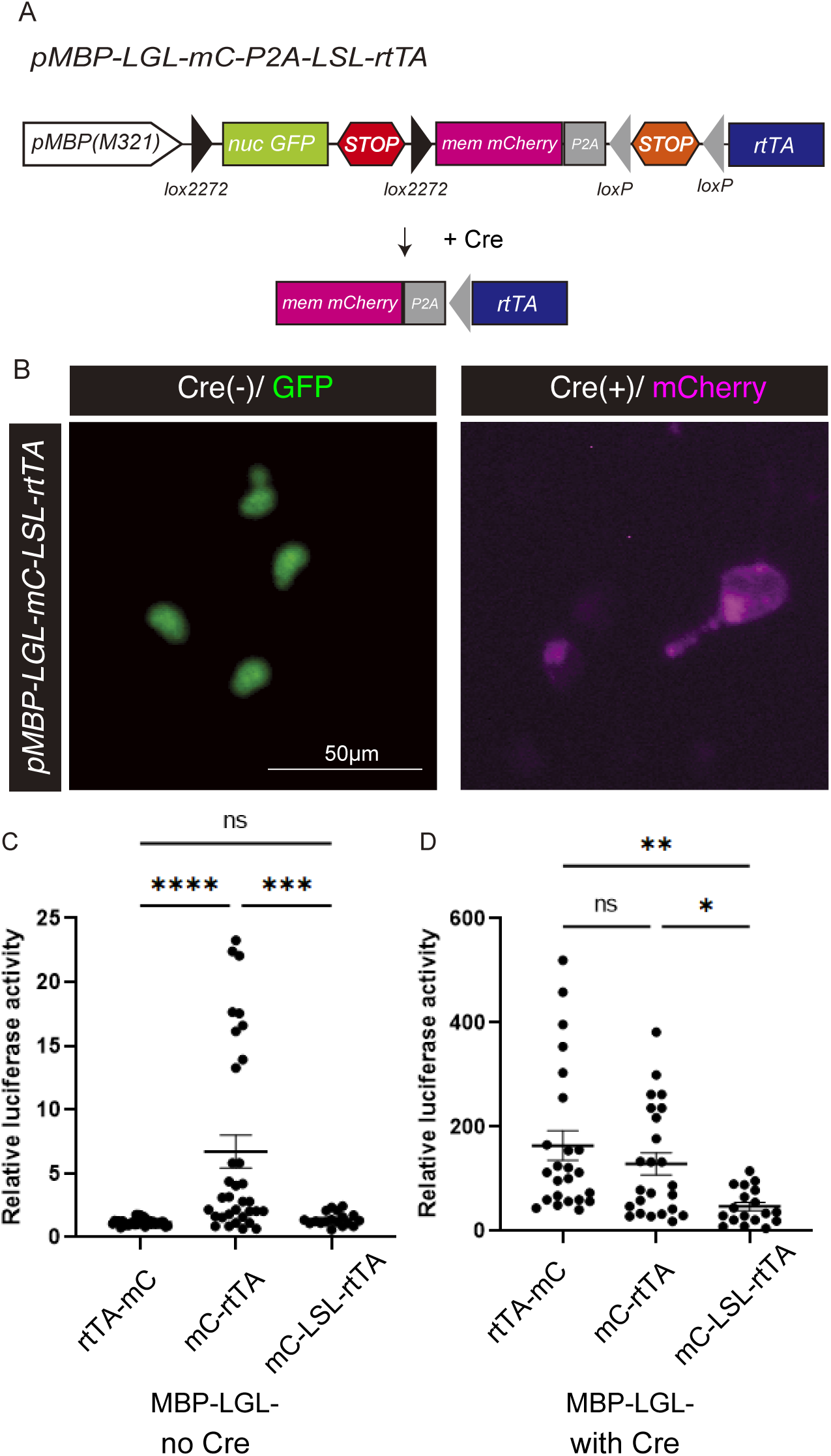
Insertion of lox-STOP-lox between P2A and 3′ transgene reduced the leaky expression of 3′ transgene. (A) Schematics diagram of *MBP-LGL-mC-LSL-rtTA*. (B) Cells transfected with the *MBP-LGL-mC-LSL-rtTA* express nucleus-targeted GFP in Cre (-) condition (left) or membrane targeted mCherry in Cre (+) condition (right). Scale bar, 50 µm. (C) Comparison of relative luciferase activity between *MBP-LGL-rtTA-mC*-, *MBP-LGL-mC-rtTA*- or *MBP-LGL-mC-LSL-rtTA*-transfected cells in Cre (-) condition (*rtTA-mC* 1.16 ± 0.08, *mC-rtTA* 11.9 ± 1.95, *mC-LSL-rtTA* 1.42 ± 0.12, n = 33, 33, 18 respectively), the data sets of *MBP-rtTA-mC* and *MBP-mC-rtTA* were obtained from Figure 2C. (D) Comparison of relative luciferase activity between cells transfected in Cre (+) (*rtTA-mC* 164.3 ± 28.7, *mC-rtTA* 129.2 ± 21.4, *mC-LSL-rtTA*47.1 ± 8.03, n = 24, 24, 18 respectively), the data sets of *MBP-rtTA-mC* and *MBP-mC-rtTA* were obtained from Figure 2D. **p< 0.01, ***p< 0.001, one-way ANOVA with Tukey test. Data shown as mean ± SEM.

### 3′ transgene leakage in the widely-used AAV-FLEX constructs

Finally, we checked whether the widely-used bicistronic inducible vectors expressed 3′ transgene leakage in vivo. We prepared three AAV-FLEX constructs from Addgene plasmids, AAV-*EF1a-FLEX-GFP-T2A-TVA* (here after *FLEX-GFP-TVA*), AAV-*βactin-FLEX-DTR-GFP* (here after *FLEX-DTR-GFP*), and AAV-*FLEX-EF1a-ArchT-GFP* (here after *FLEX-ArchT-GFP*) (**Figure 6A**). Theoretically, these vectors should express GFP only in Cre (+) condition. We stereotaxically injected a mixture of each AAV-FLEX-construct and AAV-Cre-mCherry to check whether those AAV vectors correctly express GFP in Cre (+) condition. After immunostaining for GFP, cells near the injection site were immunoreactive for GFP and mCherry in all AAV-FLEX constructs (**Figure 6B**). The ratio of GFP/Cre-mCherry-double positive cells was the lowest in *FLEX-DTR-GFP*-injected cells (*GFP-TVA* 46.4 ± 6.48 %, *DTR-GFP* 20.9 ± 3.23 %, *ArchT-GFP* 39.6 ± 2.45, n = 4; p = 0.007, Figure 6C), possibly because the *β* actin promoter activity of FLEX-DTR-GFP was lower than the EF1a promoter activity of *FLEX-GFP-TVA* and *FLEX-ArchT-GFP* (**Figure 6A**). Following this, we checked whether the AAV-FLEX vectors express leaky GFP in Cre (-) condition by co-injecting of AAV-FLEX vector and AAV-*mCherry*. We observed that compared to *FLEX-GFP-TVA* (5′ *GFP*), the ratio of GFP/mCherry-double positive cells was significantly higher in *FLEX-DTR-GFP* (*3*′ *GFP*) or *FLEX-ArchT-GFP* (*3*′ *GFP*)-injected cells (*GFP-TVA* 2.55 ± 0.85 %, *DTR-GFP* 11.7 ± 3.16 %, ArchT-GFP 22.6 ± 1.65, n = 5 or 6 hemispheres; p < 0.0001, **Figure 6D**). The position of GFP was 5′ in *FLEX-GFP-TVA* and 3′ in both *FLEX-DTR-GFP* and *FLEX-ArchT-GFP*. The results indicated that the 3′ transgene was leaky in bicistronic AAV vector-infected cells in vivo. Additionally, the data suggested that the 3′ leakage was not driven by general promoters because GFP leakage of *FLEX-DTR-GFP* (βactin promoter) was higher than that of *FLEX-GFP-TVA* (EF1a promoter) in Cre (-) condition while the expression level was opposite in Cre (+) condition.

**Figure 6.**
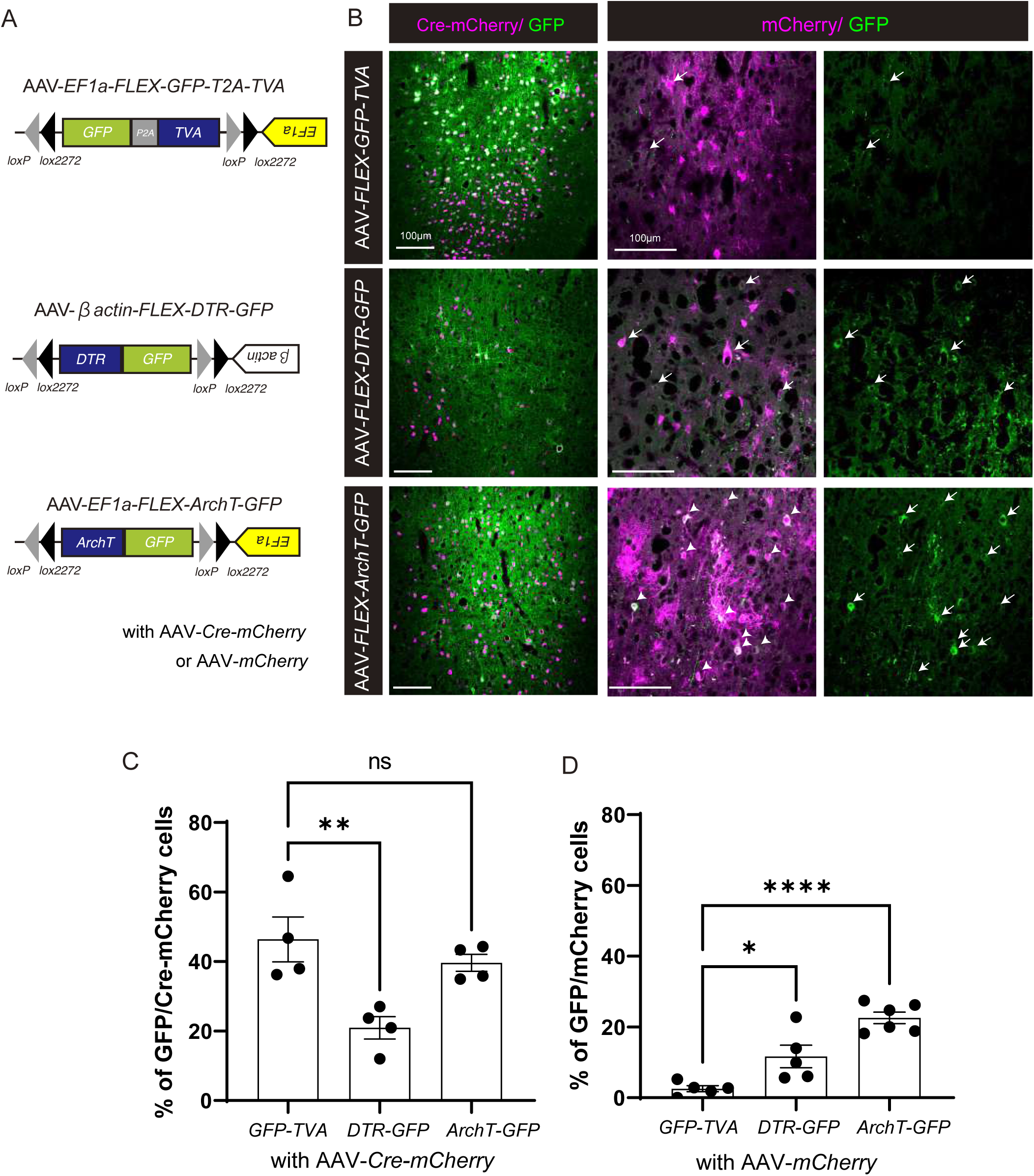
Bicistronic AAV-FLEX vectors express 3′ transgene leakage in vivo. (A) Schematic diagram of AAV-*EF1a-FLEX-GFP-T2A-TVA*, AAV-*βactin-FLEX-DTR-GFP,* and AAV-*EF1a-FLEX-ArchT-GFP*. (B) Representative images of mouse brains injected with *FLEX-GFP-TVA*, *FLEX-DTR-GFP,* or *FLEX-ArchT-GFP* and co-injected with AAV-*Cre-mCherry* or AAV-*mCherry*. Arrows indicate cells co-expressing mCherry and leaky GFP. Scale bars, 100 µm. (C) Comparison of the ratio of cells expressing GFP and Cre-mCherry between brain hemispheres injected with *FLEX-GFP-TVA*, *FLEX-DTR-GFP,* or *FLEX-ArchT-GFP*, co-injected with AAV-*Cre-mCherry* (*GFP-TVA* 46.4 ± 6.48 %, *DTR-GFP* 20.9 ± 3.23 %, *ArchT-GFP* 39.6 ± 2.45, n = 4 hemispheres). (D) Comparison of cells expressing leaky GFP and mCherry between brain hemispheres injected with *FLEX-GFP-TVA*, *FLEX-DTR-GFP,* or *FLEX-ArchT-GFP*, co-injected with AAV-*mCherry* (*GFP-TVA* 2.55 ± 0.85 %, *DTR-GFP* 11.7 ± 3.16 %, *ArchT-GFP* 22.6 ± 1.65, n = 5 or 6 hemispheres). n.s. p > 0.05, *p < 0.05, **p < 0.01, ****p < 0.0001, one-way ANOVA with Dunnett’s test. Data shown as mean ± SEM.

## Discussion

In this study, we observed that coding sequences located in the 3′ region of polycistronic expression vectors were leaky, and they were not derived from normal promoter activity. Thus, our data indicated that using a polycistronic expression strategy such as fusion protein, including IRES or 2A peptide, better not to be combined with an inducible gene expression strategy such as lox-STOP-lox, FLEX, TetOn/Off, or cell-specific promoters, especially in transient expression experiments.

Some reports have indicated leaky expression in an inducible gene expression strategy (Lindeberg and Ebendal, 1999) (Luche et al., 2007) (Igarashi et al., 2016) (Lavin et al., 2020). However, this is often ignored by researchers if results of the experiment are unaffected by the leaky expression. For example, if the expression level of leaky GFP or any other fluorescent protein is low enough, the signal will not be detected and data analysis will not be affected. Yet, we observed that more than 7% of the cells expressed mCherry without Cre activity in *pCAG-LGL-rtTA-P2A-mC*-transfected cells. Noteworthy, we did not use any antibody to observe leaky mCherry expression, and the fluorescence was detected without signal enhancement. Thus, analysis of cells expressing the 5′ CDS gene in bicistronic vector could be confused with leaky 3′ fluorescent gene expression. Additionally, even a few molecules of some proteins can execute their functions. For example, DTA is a widely used suicide gene (Lee et al., 1998) (Stanger et al., 2007), and one molecule of DTA is sufficient to kill the cell (Yamaizumi et al., 1978). Our results suggested that such genes should be located at the 5′ region of bicistronic vector because 5′ CDS gene expression is finely regulated as compared to 3′ CDS gene expression in bicistronic vector with inducible gene expression systems.

Not only mCherry and rtTA, in all tested CDSs, including GFP, TFP, DTR, and ArchT could also drive leaky expression. All tested bicistronic Cre-inducible vectors showed the leaky expression, including the IRES vector and the DIO vector (**Figure S1C-S1E**). Thus, it is likely that all bicistronic gene expression vectors with an inducible gene expression system have a risk of leaky 3′ CDS expression.

We tried to reduce the leaky expression by removing TF binding motifs. However, contrary to our expectations, the leaky expression increased (**Figure S2**). Results of segment transfection experiments with *rtTA*, *P2A-rtTA,* and *mC-P2A-rtTA* segments clearly showed that the 5′ region CDS was responsible for leaky rtTA expression. It is still not clear how the 5′ region CDS drove the expression of 3′ region CDS. It is widely known that RNA polymerase II should bind to the promoter DNA to initiate CDS transcription (Cramer, 2019). One possibility is that all protein CDS possess DNA sequences that can bind to RNA polymerase II, and 5′ CDS in the bicistronic construct drives the leaky expression of 3′ CDS. As demonstrated by the Callaway group (Fischer et al., 2019), removing the ATG start codon in the 5′ CDS may reduce the leakage, however, it is not feasible to remove all ATGs within a CDS since methionine is only encoded by the ATG codon and removal of all ATGs in an attempt to prevent any downstream expression, particularly in-frame ATGs, may cause loss of protein function.

We successfully reduced leaky gene expression by integrating the bicistronic construct into the genome or by inserting the lox-STOP-lox cassette between the P2A sequence and 3′ CDS region (**Figure 5**). In *MBP-LGL-mC-P2A-LSL-rtTA*-transfected cells, mCherry and rtTA expression was tightly regulated by Cre recombination. However, we found that rtTA expression level following Cre recombination was lower than that of normal constructs. Following are some possible explanations for the low rtTA expression in *MBP-LGL-mC-P2A-LSL-rtTA*-transfected cells. 1) recombination efficiency; both *lox2272-nG-STOP-lox2272* and *loxP-STOP-loxP* were used in the construct. It might be possible that only one of lox-STOP-lox cassettes was removed but the other cassette was not removed. 2) additional peptides at the N-terminal of rtTA protein; loxP DNA segment was present between P2A and rtTA CDS following Cre recombination, and transcription from loxP sequence added extra peptides at the N-terminal of rtTA protein. This could be the cause of reduced rtTA activity. Although the expression level of rtTA following Cre recombination was low, this construct was useful because the expression levels of leaky mC and leaky rtTA were minimum (**Figure 5**).

The simplest method to express two genes under the control of a specific promoter is by using two separate constructs with an identical promoter. We generated a plasmid with two MBP promoters having different orientations. Cells transfected with the construct successfully expressed two different genes without leaky gene expression (data not shown). Some bidirectional promoter constructs such as MXS_BidirectionalCAG are available from Addgene (Sladitschek and Neveu, 2016) (Addgene #78175). Such bidirectional vectors would be safer than using a bicistronic vector with an inducible gene expression system.

It has been reported that *somatostatin-IRES-Cre* mouse expresses Cre in non-somatostatin-expressing neurons (Hu et al., 2013). It is likely that the leaky Cre expression was driven by somatostatin CDS, similar to our result. We contacted the inventor of the mouse line possessing *CAG-lox-STOP-lox-RABVgp4-IRES-TVA* gene (Takatoh et al., 2013). The author kindly informed us that there were signs of leaky TVA expression while RABVgp4 was correctly regulated by Cre recombination (personal communication). Despite these limitations, the mouse line can still be used to target the peripheral nervous system (Zhang et al., 2015). Some studies have reported no leaky expression of genes in the bicistronic vector with an inducible gene expression system (Atasoy et al., 2008; Yamamoto et al., 2009). Atasoy and colleagues generated *AAV-FLEX-ChR2-mCherry* (2008), a useful tool for investigating the role of a particular neuronal subtype. The authors reported no leaky mCherry expression; however, if you carefully observe the figures with changing scale and contrast, leaky mCherry expression was observed. It is likely that the criteria for counting mCherry-positive cells were high for the study.

Taken together, this study provided an important caveat to using bicistronic vector with inducible gene expression. We recommend using two separate vectors with an identical promoter to induce the expression of two transgenes or using a vector containing bi-directional promoter. If you need to use a bicistronic vector to induce the expression of a functional gene, such as ArchT and DTR, plus a fluorescent reporter gene such as GFP, it is better to put a functional gene on 5′ region CDS and a fluorescent reporter gene on the 3′ region because functional genes such as ArchT and DTR significantly affect the experimental data as compared to the leaky expression of fluorescent reporter genes. This finding is important to generate constructs to precisely control multiple gene expression for basic studies and gene therapy.

## Acknowledgements

This work was supported by The Naito Foundation Subsidy for Inter-Institute Researches (Y.O.), KAKENHI Grants from the Japan Society for the Promotion of Science (20K22691 & 21K15197, Y.O.; 21H05241, 21H04786, 20K21506, & 20KK0170, N.O.), a research grant from the National Center of Neurology and Psychiatry (No. 30-5, 3-5; N.O.), a Future Fellowship from the Australian Research Council (FT150100207, T.M.), and generous philanthropic support provided by Metal Manufactures Ltd. (T.M.). The Australian Regenerative Medicine Institute at Monash University is supported by grants from the State Government of Victoria and the Australian Government. We gratefully acknowledge support provided by research platforms at Monash University, including: FlowCore for flow cytometry and fluorescence-activated cell sorting (FACS), Micromon for Sanger sequencing and Monash Micro Imaging for light microscopy. We thank the following researchers for plasmids deposited at Addgene that we have used in this study: Geoff Wahl for *H2B-GFP* (Addgene #11680), Liqun Luo for *pCA-mTmG* (Addgene #26123), Connie Cepko for *pCAG-Cre* (Addgene #13775), Joseph Loturco for *PBCAG-mRFP* and *pCAGPBase* (Addene #40996, #40972), Michael Davidson for *mTFP1-H2A-10* (Addgene #55488), Hongkui Zeng for the *Ai65(RCFL-tdT)* targeting vector (Addgene #61577), Edward Callaway for *pAAV-EF1a-FLEX-GT* (Addgene #26198), Eiman Azim and Thomas Jessell for *pAAV-FLEX-DTR-GFP* (Addgene #124364) and Edward Boyden for *pAAV-EF1a-FLEX-ArchT-GFP* (Addgene #58851). We also thank Jun Takatoh for providing important information about R Φ GT mouse line and Mirana Ramialison (ARMI) for advice on approaches to identify transcription factor binding motifs.

## Materials and Methods

### Molecular cloning

Plasmid vectors were constructed using restriction cloning, PCR cloning, and Gibson cloning (Gibson et al., 2009). Some DNA segments were synthesized using gBlock gene fragments (Integrated DNA Technologies, USA). Details of the cloning steps with primer sequences are described in *Supplemental Information*. DNA segments obtained by restriction digestion or PCR amplification were purified using the QIAquick spin column (28115; QIAGEN, Germany). All plasmid constructs were verified by Sanger sequencing at Micromon, Monash University.

### Gel extraction

PCR solution or enzymatically digested plasmid solution was purified via agarose gel extraction. Agarose gel (1 %) was prepared in 1 x TAE and SYBR safe solution (Thermofisher). DNA solution was added to the 6X loading buffer and a total of 120-180 µL of solution was loaded into a combined well, where three wells of a gel maker were combined to one well (AD216, Clonetech). Gels were run at 130 V for 45 min in an electrophoresis apparatus (Horizon11.14, Thermofisher). Bands of correct size were excised and purified using QIAquick gel extraction kit (Qiagen). The DNA segments used for cell transfection experiments were further purified by Phenol/Chloroform extraction and ethanol purification.

### Cell culture

HEK293T cells were cultured in Dulbecco’s Modified Eagle Medium (DMEM; Invitrogen, USA) supplemented with 10% fetal bovine serum (FBS; Corning, USA) and 1% Penicillin-Streptomycin-Glutamine (Gibco, USA). CG4 cells (Louis et al., 1992) were cultured in SATO medium supplemented with N-Acetyl-Cysteine (60 µg/mL, Sigma), biotin (10 ng/mL, Sigma), forskolin (5 µM, Sigma), Insulin (5 µg/mL, Sigma), recombinant human NT3 (5 ng/mL, Peprotech) and PDGF-AA (10 ng/mL, Peprotech). SATO medium was prepared in high-glucose DMEM (Gibco, Cat. 31966-021) using 100x SATO stock solution which was prepared as described in Cold Spring Harbor Protocols (doi:10.1101/pdb.rec077073).

### Transfection

Transfections were performed one day after seeding cells into 96-well plates (1.0 x 10^4^ HEK293T cells/well; 0.9 x 10^4^ CG4 cells/well) or into 24-well plates (1.0 x 10^5^ HEK293T cells/well; 5.0 x 10^4^ CG4 cells/well) using Lipofectamine LTX reagent with PLUS reagent (Thermofisher, USA) in medium without Penicillin-Streptomycin according to the manufacturer’s protocols.

For HEK 293T cells, plasmid DNA (0.5 µg per construct) and PLUS reagent (0.5 µL per construct) were diluted in 50 µL Opti-mem (Invitrogen). This solution was mixed with Opti-mem containing Lipofectamine LTX (1 µL Lipofectamine LTX in 50 µL Opti-mem, incubated for 5 min). After 20 min of incubation at room temperature, the transfection solution was added to each well (100 µL/well for 24 well plate; 20 µL/well for 96-well plate). The medium was replaced with 10% FBS-DMEM medium containing Penicillin-Streptomycin 4 h post transfection.

For CG4 cells, the cells were incubated in SATO medium containing PDGF-AA and NT3 during transfection. Constructs (each 0.5 µg) and PLUS reagent (0.5 µL for one construct) were diluted in 25 µL Opti-mem (Invitrogen). The Opti-mem containing constructs and PLUS reagents was mixed with the Opti-mem/ Lipofectamine LTX (2 µL Lipofectamine LTX in 25 µL Opti-mem pre-incubated for 5 min). After 20 min of incubation at room temperature, the mixed solution was added to each well (50 µL/well for 24 well plate; 10 µL/well for 96-well plate). The medium was replaced to SATO medium containing T3 (40ng/mL, Sigma) and Penicillin-Streptomycin 4 h post transfection.

### Luciferase assays

The pTRE-Luciferase (Clontech) and constructs containing rtTA transgene were co-transfected into HEK 293T or CG4 cells using Lipofectamine LTX. Four hours after Lipofectamine transfections, the medium was replaced with medium containing doxycycline (1 µg/mL) and Penicillin-Streptomycin. T3 (40 ng/mL) was also added to the medium to induce differentiation of CG4 cells into oligodendrocytes. Two days post transfection, luciferase assays were performed using a luciferase assay kit (Pierce) according to manufacturer’s protocol. The average chemiluminescence intensity for wells transfected with pTRE-Luciferase alone was defined as having a relative luciferase activity (RLA) of one.

### Flow cytometry

Flow cytometry was performed on a BD Fortessa™ X-20 cell analyser with BD FACSDiva software (BD Biosciences, USA). The voltage, compensation, scale and gating were determined by first adjusting them for negative and single fluorophore-only controls. Fluorophore Minus One (FMO) controls (DAPI + mCherry, DAPI + GFP, and GFP + mCherry samples) were also used for initial adjustment of compensation. For each sample, 10,000 events were acquired. Data were first gated to include only intact cells (FSC-A vs. SSC-A) and then single cells (FSC-A vs. FSC-H), and DAPI-negative cells were gated as live cells (FSC-A vs. BV450-A). Fluorophore expression was observed in live cells. Once the parameters were established, the same settings and gate parameters were used to acquire and analyse all other samples. The acquired data were analysed using FlowJo version 10 (BD Biosciences), and the same compensation, scale, and gating parameters were used for each experiment. For each experiment, a plot comparing the proportion of GFP-expressing cells vs. the proportion of mCherry-expressing cells was generated, and these plots are presented in *Results*

### Generation of stable cell lines

HEK-*CAG-LGL-rtTA-P2A-mC* or HEK-*CAG-LGL-mC-P2A-rtTA* stable cell lines were generated using the PiggyBac transposon-based expression system (Li et al., 2013). Briefly, the *PB-CAG-LGL-rtTA-P2A-mC* or *PB-CAG-LGL-mC-P2A-rtTA* construct was co-transfected with *pCAGPBase* (Addgene #40972) into HEK293T cells. The transfected cells were passaged every 2-4 days for 20 days. Following this, FACS sorting was performed to collect the top 10% of nGFP-expressing cells. The collected nGFP (+) cells were expanded and used as HEK*-CAG-LGL-rtTA-P2A-mC* or HEK*-CAG-LGL-mC-P2A-rtTA* stable cell lines.

### Mutagenesis of putative TF binding motifs

Putative transcription factor (TF) binding motifs within the mCherry CDS were predicted using the JASPAR 2018 database (Khan et al., 2018). We selected putative TF binding sites using the following criteria: 1) JASPAR score > 0.9, 2) defined as a transcriptional activators (transcriptional repressors were ignored), 3) only TFs expressed in oligodendroglia were included by cross-referencing against the Brain RNA-Seq website ((Zhang et al., 2014); https://web.stanford.edu/group/barres_lab/brain_rnaseq.html). Next, we defined a TF-modified version of the membrane-targted mCherry CDS by incorporating synonymous mutations that altered the DNA sequence without affecting the amino acid sequence in order to mutate the selection of putative TF binding sites (*see Supplementary Information*).

### Animals

Wild type 8-week-old C57BL6 female mice were purchased from Clea Japan or Japan SLC. Mice were housed in a 12-hour light/dark cycle with free access to water and food. Animal experiments were conducted according to the guidelines of Jichi Medical University.

### AAV preparation and injection

AAV2-*EF1a-FLEX-GFP-T2A-TVA*, AAV2-*βactin-FLEX-DTR-GFP* and AAV2*-EF1a-FLEX-ArchT-GFP* were generated from pAAV-*EF1a-FLEX-GT* (Addgene #26198) (Wall et al., 2010), *pAAV-FLEX-DTR-GFP* (Addgene #124364), and pAAV-*EF1a-FLEX-ArchT-GFP* (Addgene# 58851) (Han et al., 2011), respectively, as previously described (Watakabe et al., 2015) (Osanai et al., 2017). AAV1-*Cre-mCherry* and AAV2-*mCherry* were purchased from SignaGen laboratory (SL101117 and SL101405, respectively). Either AAV1-*Cre-mCherry* or AAV2-*mCherry* and one from AAV2-*EF1a-FLEX-GFP-T2A-TVA*, AAV2-*βactin-FLEX-DTR-GFP,* and AAV2*-EF1a-FLEX-ArchT-GFP* were mixed (1.0×10^12^vg/mL for each AAV) for stereotaxic injection.

Mice were anaesthetized with a cocktail of medetomidine (0.3 mg/kg), midazolam (4.0 mg/kg) and butorphanol (5.0 mg/kg) as described previously (Yamazaki et al., 2021). Mice were placed in a stereotaxic frame (Narishige) with a mouse adaptor. After drilling a hole into the skull to expose the injection site, 1 μL (1.0×10^9^vg) of AAV solution was stereotaxically injected into the somatosensory cortex (1.0 mm posterior and 1.5 mm lateral to the bregma, at a depth of 0.3 mm) using NanojectII as described previously (Osanai et al., 2017). Mice were sacrificed 14 days after AAV injection.

### Tissue processing and immunohistochemistry

Mice were perfused transcardially with 4% paraformaldehyde (PFA) in 0.1 M sodium phosphate buffer (pH 7.4) under the deep anesthesia. The brains were collected from each animal and postfixed with 4% PFA overnight at 4 ℃. Postfixed brains were cryoprotected overnight in 0.1 M sodium phosphate buffer (pH 7.4) containing 30% sucrose, embedded in OCT compound (Sakura Finetek), and cut into 10 µm sections using a cryostat (CM3050; Leica). The sections were blocked in 10% normal goat serum in D-PBS with 0.3% Triton X-100 (PBST) then incubated with rat anti-GFP antibody (1:1000; GF090R; Nacalai Tesque) in PBST at 4 ℃ overnight. After washing with PBST, the sections were incubated with secondary antibodies (1:500, Alexa Fluor 488- conjugated goat anti-rabbit IgG; Molecular Probes) for 1 h at room temperature. Slides were rinsed again and counterstained with Hoechst 33342 (1 µg/mL; Invitrogen), and coverslipped as described previously(Oluich et al., 2012).

### Microscopic analysis

The number of GFP (+) and/or mCherry (+) cells in the wells of 96-well plate was counted manually using the 10× objective of an Olympus IX81 inverted fluorescence microscope (Olympus). All cells in the well were counted.

Sections from AAV-injected brains were examined under 20x objective on an FV1000 confocal microscope (Olympus). Cell counts of confocal images were performed using Fiji software. Images of Hoechst, GFP, and mCherry were changed to binary images through Fiji threshold function. The binary images of Hoechst, GFP, and mCherry were merged, and co-localization of signals was analysed manually. Fluorescent signals exhibiting clear cellular morphology were included in the cell quantified.

### Statistical analysis

Prism 7 (GraphPad) and Excel (Microsoft) were used for statistical analysis. Cell counting experiments and luciferase assays were repeated at least three independent experiments. The statistical significance level was set at *p* < 0.05. All data are described as mean ± SEM.

