## Supplemental information for "5′ transgenes drive leaky expression of 3′ transgenes in inducible bicistronic vectors"

Supplementaly figure1

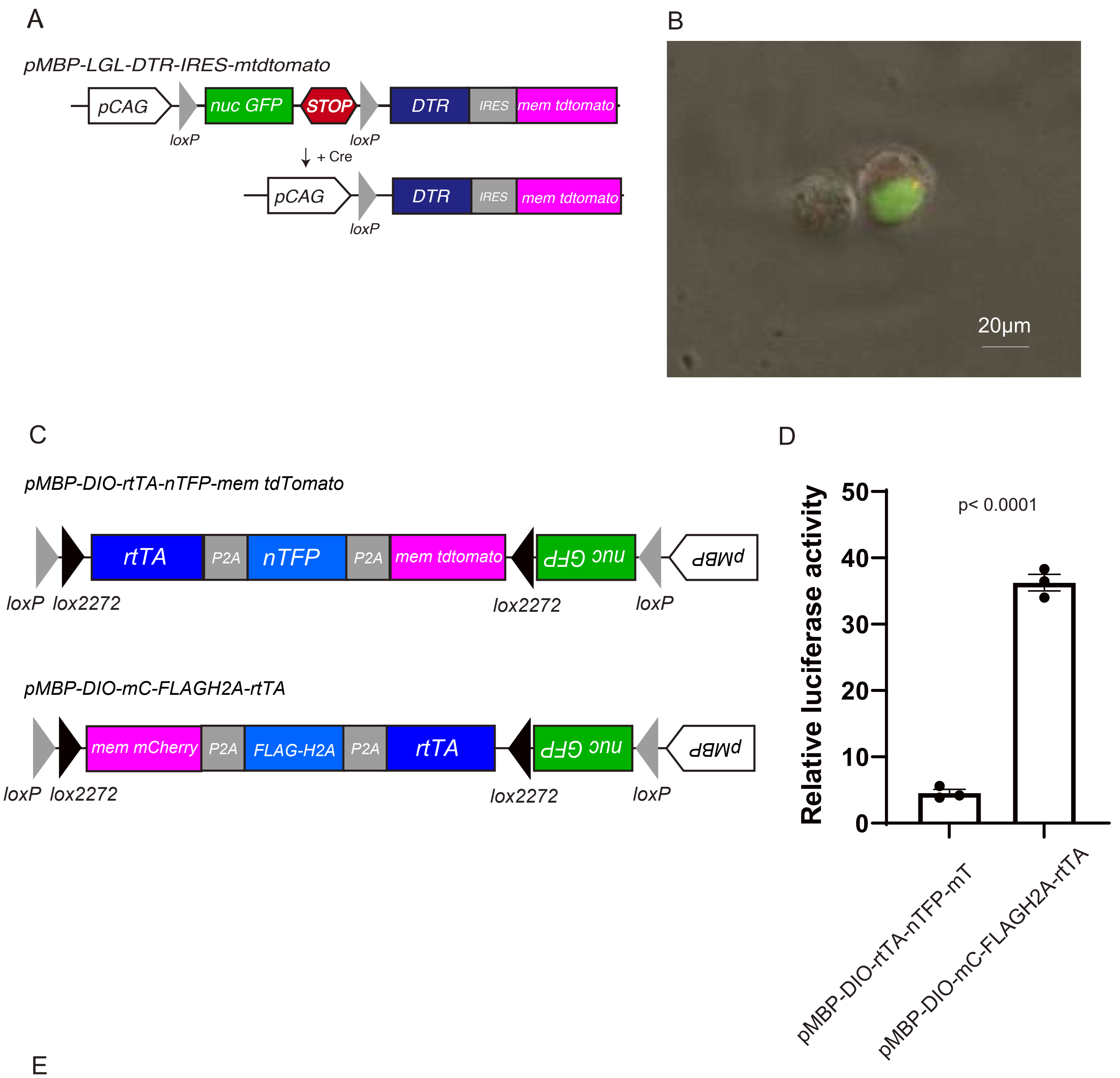


E

| | # of GFP (+) | # of nGFP/mT or mC double positive cells | % of mT or mC positive cells/ GFP positive cells |
|---|---|---|---|
| *pMBP-LGL-tdTomato* | 127 | 0 | 0.0 |
| *pMBP-LGL-DTR-IRES-mT* | 101 | 9 | 8.9 |
| *pMBP-LGL-DTR-P2A-TFP-P2A-mT* | 50 | 8 | 16.0 |
| *pMBP-LGL-mC-DTR* | 312 | 0 | 0.0 |
| *pMBP-LGL-mC-FLAG-H2A-rtTa* | 329 | 0 | 0.0 |

### Supplemental figure2

A

| Normal mCherry | Modified mCherry |
|---|---|
| SPI1 TF binding motif | No TF binding motif |
| … ATG GGA AGC … | … ATG GGT AGC … |
| … Met Gly Ser … | … Met Gly Ser … |

C

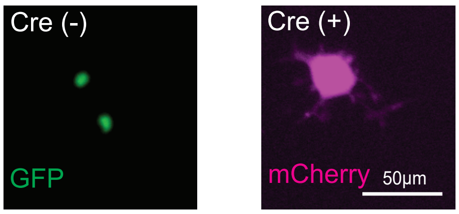


B

| Gene ID | Name | Score | Sequence | Gene expression (FPKM) |
|---|---|---|---|---|
| MA0080.1 | SPI1 | 0.99446381 | gggaag | 11.06 |
| MA0765.1 | ETV5 | 0.95528969 | agcggatgta | 34.82 |
| MA0512.1 | Rxra | 0.94860762 | ctgaggtcaag | 7.4 |
| MA0830.1 | TCF4 | 0.93963159 | tgcagctgcc | 50.7 |
| MA0831.2 | TFE3 | 0.93869785 | cgcgtgat | NA |
| MA0671.1 | NFIX | 0.93592514 | accgccaag | 22.48 |
| MA0018.2 | CREB1 | 0.93420224 | tgaggtca | 5.99 |
| MA0259.1 | ARNT::HIF1A | 0.93257452 | ctacgtga | 36.32 |
| MA0764.1 | ETV4 | 0.93082669 | agcggatgta | 54.69 |
| MA0522.2 | TCF3 | 0.92975001 | tgcagctgcc | 24.35 |
| MA0671.1 | NFIX | 0.92747045 | aaggccaag | 22.48 |
| MA0761.1 | ETV1 | 0.92508093 | agcggatgta | 13.76 |
| MA0614.1 | Foxj2 | 0.92377284 | gtcaacat | 8.05 |
| MA1100.1 | ASCL1 | 0.92137301 | cgtgcagctgcc | 9.85 |
| MA0599.1 | KLF5 | 0.91956037 | actcctccct | 0.32 |
| MA0522.1 | Tcf3 | 0.91757382 | tgcagctgccc | 24.35 |
| MA0727.1 | NR3C2 | 0.91549275 | tgggacatcctg | 2.32 |
| MA0259.1 | ARNT::HIF1A | 0.90302713 | gcgcgtga | 36.32 |
| MA0827.1 | OLIG3 | 0.90236088 | agcatatggg | 0.1 |
| MA0826.1 | OLIG1 | 0.90208241 | agcatatggg | 1359.97 |

D

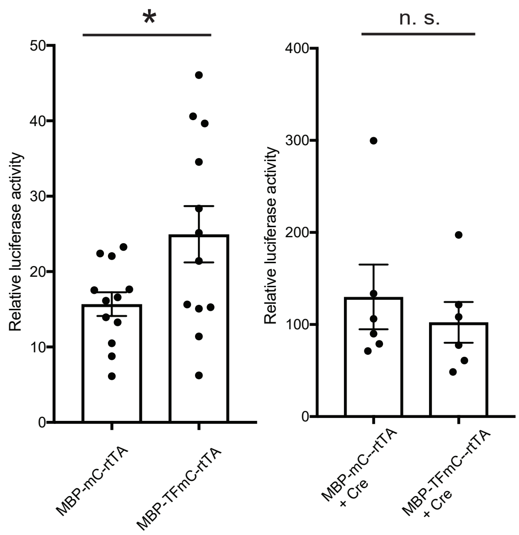

**Supplementary Figures**

**Figure S1. Leaky 3′ transgene expression derived by several types of 5′ transgenes in IRES, DIO vectors.**

**(A)** Scheme of *pMBP-LGL-DTR-IRES-mem tdtomato*. **(B)** Representative picture of a CG4 cell expressing leaky tdtomato. In this construct, *DTR* seems to drive *tdtomato* expression. Scale bar: 20 µm. **(C)** Scheme of *pMBP-DIO-rtTA-nTFP-mem tdtomato* or *pMBP-DIO-mC-FLAGH2A-rtTA*. **(D)** Relative luciferase activity of CG4 cells that were transfected with *pMBP-DIO-rtTA-nTFP-mem tdtomato* plus *pTRE-Luciferase* or *pMBP-DIO-mC-FLAGH2A-rtTA* plus p*TRE-Luciferase* in Cre (-) condition. *pMBP-DIO-mC-FLAGH2A-rtTA* plus *pTRE-Luciferase* transfected CG4 cells expressed higher amount of rtTA than that of *pMBP-DIO-rtTA-nTFP-mem tdtomato,* likely because *rtTA* is located at 3′ position in *pMBP-DIO- mC-FLAGH2A-rtTA*. **(E)** Table indicating % of cells expressing leaky tdTomato or mCherry in several inducible bicistronic constructs and *pMBP-LGL-tdtomato* construct. Cells transfected with *pMBP-LGL-tdtomato* construct did not express leaky tdtomato. These results suggest that *DTR* and/or *TFP* drive leaky tdtomato or mCherry expression. These data also suggest that inducible bicistronic vector using IRES sequence also expressed leaky 3′ transgene. Student’s t-test $p < 0.0001$. Data shown as mean ± SEM.

**Figure S2. Modification of 5′ transgene DNA sequence altered 3′ transgene expression in inducible bicistronic vectors.**

**(A)** Scheme of a strategy to remove transcription factor (TF) binding site. **(B)** Table showing TF binding sites in the coding sequence of membrane-targeted mCherry. The TF binding site data were obtained from JASPAR 2018 database (Khan et al., 2018), and data of TF expression level in murine oligodendrocyte progenitor cells (OPCs) were obtained by Brain

RNA-Seq website. Notably the CG4 cell line was derived from rat OPCs. **(C)** Representative images of cells transfected with *pMBP-TF modified mC-P2A-rtTA* in Cre negative condition (left) or positive (right) condition. Transfected cells correctly expressed fluorescent proteins in Cre-dependent manner. Scale bar 50 μm. **(D)** Relative luciferase activity of cells transfected with normal *pMBP-mC-P2A-rtTA* or *pMBP-TF modified mC-P2A-rtTA* in Cre negative (left) or positive (right) condition. Unexpectedly, cells transfected with the TF removed *pMBP* construct expressed higher amount of rtTA compared to normal *pMBP-mC-P2A-rtTA* transfected cells, while both the constructs expressed similar amount of rtTA in Cre (+) condition. Student t-test n.s. $p > 0.05$, *$p < 0.05$. Data shown as mean ± SEM.

### Supplementary Methods

#### Generation of plasmid constructs

##### *(i) pMBP-loxP-nG-STOP-loxP*

*pMBP-DTR* (Oluich et al., 2012) was enzymatically digested by *BamHI,* and pMBP backbone was collected by gel extraction using a gel extraction kit (#28704, QIAGEN) and re-ligated to obtain *pMBP* backbone. The *loxP-H2B-GFP* fragment was PCR amplified using *H2B-GFP* plasmid (Addgene# 11680) (Kanda et al., 1998) and following primers:

*Sal1-loxP-H2B-EGFP-Xho1-Sal* F: (5′-GCTAGTCGACATAACTTCGTATAGCATACATTATACGAAGTTATGCCACCATGCCAGAGCCAGCGA-3′)

*Sal1-loxP-H2B-EGFP-Xho1-Sal* R: (5′-TAGCGTCGACCTCGAGTTACTTGTACAGCTCGTCCATG-3′). The *loxP-H2B-GFP* fragment was inserted into *SalI* site of *pMBP*. As a result, *pMBP-loxP-nG* plasmid was obtained. To generate *pMBP-loxP-nG*-STOP-*loxP* plasmid, STOP-*loxP* fragment was inserted into *XhoI* site of *pMBP-loxP-nG* construct. The STOP-*loxP* fragment was PCR amplified using *pCA-mTmG* plasmid (Addgene # 26123) and following primers:

*STOP-loxP* F: (5′-GCTACTCGAG GGATCTTTGTGAAGGA-3′)

*STOP-loxP* R: (5′-TAGCCTCGAGTCTAGATAGCCATATGGGTACCGATATCATAACTTCGTATAATGTATGC-3′).

##### *(ii) pMBP-loxP-nG-STOP-loxP-rtTA-P2A-TFP*

*rtTA-P2A-TFP* fragment was generated by combining *rtTA-5′ P2A* fragment and *3′ P2A- TFP* fragment through PCR as following.

*rtTa-5′ P2A* fragment was generated by PCR using *pCMV-Tet3G* (Clontech) as template and *rtTa-P2A* FW primer (5′-GCTAGATATCTTCACCATGTCTAGACTGGAC-3′) and *rtTa-P2A* RV primer (5′-AAGTTAGTAGCTCCGCTTCCTCCTGGTAACATGTCAAGGTCAAAATCGTCAA-3′).

*3′ P2A- TFP* fragment was generated by PCR using *mTFP1-H2A-10* as template, which was a gift from Michael Davidson (Addgene # 55488) (Ai et al., 2008), and using *P2A-TFP* forward primer (5′-CAGCCTGCTGAAGCAGGCTGGAGACGTGGAGGAGAACCCTGGACCTATGTCGGGACGTGGCAAGCAGGG-3′) and *P2A-TFP* reverse primer (5′-TAGCGGTACCCTTGTACAGCTCGTCCATGCCG-3′). PCR solution containing *rtTA-5′ P2A* fragment and that containing *3′ P2A- TFP* fragment were 1:1 mixed and 2 μL of the mixed solution was used as template in total 50 μL of PCR solution, and the *rtTA-P2A* forwrd primer and the *P2A-TFP* reverse primer were used to generate *rtTA-P2A-TFP* fragment. PCR amplified *rtTA-P2A-TFP* fragment was inserted into *EcoRI – KpnI* site of *pMBP-loxP-nG-STOP-loxP* construct.

***(iii) pMBP-loxP-nG-STOP-loxP-mC-P2A-DTR***

*EcoRV-Membrane mCherry* (hereafter mC) *-P2A-KpnI-DTR-KpnI-XbaI* fragment was synthesized (gBlocks Gene Fragment, IDT) and PCR amplified. The fragment was digested by *EcoRV* and *XbaI* and inserted into *EcoRV-XbaI* site of *pMBP-loxP-nG-STOP-loxP plasmid.*

***(iv) pMBP-loxP-nG-STOP-loxP-mC-P2A-FLAG-H2A-P2A-rtTa-WPRE***

*pMBP-loxP-nG-STOP-loxP-mC-P2A-DTR* construct was digested with *KpnI* and *pMBP-loxP-nG-STOP-loxP-mC-P2A* fragment was collected by gel extraction. *KpnI-FLAG-H2A-P2A-rtTa-WPRE-KpnI* fragment was synthesized (gBlocks Gene Fragment, IDT) and PCR amplified and digested by *KpnI*. *pMBP-loxP-nG-STOP-loxP-membrane mCherry-P2A* fragment and *KpnI-FLAG-H2A-P2A-rtTa-WPRE-KpnI* fragment were ligated with *KpnI* site.

***(v) pMBP-loxP-nG-STOP-lox2272-EcoRV-loxP-lox2272***

*STOP-lox2272* fragment was generated by PCR amplification using *pMBP-loxP-nG-STOP-loxP* as a template and *STOP-lox2272* forward primer (5′- GCTACTCGAG GGATCTTTGTGAAGGA-3′) and *STOP-lox2272* reverse primer (5′-tagcctcgagtctagatagccatatgataacttcgtataaagtatccTATACGAAGTTATTAGGTCCCT-3′). The STOP-*lox2272* fragment was inserted into *pMBP-loxP-nG* plasmid using *XhoI* site and *pMBP-loxP-nG-STOP-lox2272* was generated. The *NdeI-EcoRV-loxP-lox2272-XbaI* fragment was synthetized (gBlock, IDT) and PCR amplified. The segment was inserted into the *pMBP-loxP-nG-STOP-lox2272* plasmid digested with *NdeI* and *XbaI*.

***(vi) pCAG-loxP-nG-STOP-loxP-rtTA-P2A-mC***

Isothermal assembly technique was used for making this construct. *pCAG-Cre* (Addgene plasmid # 13775) (Matsuda and Cepko, 2007) was digested with *EcoRI* and *NotI*. The *pCAG* backbone was collected by gel extraction. The *loxP-nG-STOP-loxP-rtTA* fragment was obtained by PCR amplification using *pMBP-loxP-nG-STOP-loxP-rtTA-P2A-TFP* plasmid as template. *TPC1* forward primer (5′-GCTGTCTCATCATTTTGGCAAAGCGAATAATTCTAGGGTCGAC – 3′) and *TPC1* reverse primer (5′-TCCGCTTCCTCCTGGTAACATGT-3′), *P2A* fragment was amplified by PCR using synthetic *P2A-mtdTomato* plasmid (GeneArt, Invitrogen) as a template and *TPC2* forward primer (5′-

ACATGTTACCAGGAGGAAGCGGAGGAAGCGGAGCTACTAACTTCAG -3′) and *TPC2* reverse (5′ – GGTCTTGGAGAAACAGCAACCCAT – 3′) primer, *mC* fragment was PCR amplified using *pMBP-loxP-nG-STOP-mC-P2A-DTR* plasmid as a template and *TPC3* forward primer (5′- ATGGGTTGCTGTTTCTCCAAGACC – 3′) and *TPC3* reverse primer (5′ – GGCAGCCTGCACCTGAGGAGTGCTTATCCGCTTCCCATATGCTTGTA – 3′). These *pCAG* fragment, *loxP-nG-STOP-loxP-rtTA* fragment, *P2A* fragment and *mC* fragment were purified by gel extraction and assembled in an isothermal Gibson assembly reaction (Gibson Assembly®, New England Biolabs), followed by transformation into NEB 5-alpha competent *E. coli* (New England Biolabs). The DNA sequence of the *pCAG-loxP-nG-STOP-loxP-rtTA-P2A-mC* plasmid was confirmed by sanger sequencing.

***(vii) pCAG-loxP-nG-STOP-loxP-mC-P2A-rtTa***

This construct was generated by an isothermal Gibson assembly reaction of *pCAG* fragment, *loxP-nG-STOP-loxP-mC-P2A* fragment and *rtTA* fragment. The *pCAG* fragment was obtained by *EcoRI-NotI* digestion of *pCMV-Cre* as described above. The *loxP-nG-STOP-loxP-mC-P2A* fragment was generated through PCR amplification using *pMBP-loxP-nG-STOP-loxP-mC-P2A-FLAG-H2A-P2A-rtTA-WPRE* plasmid as a template and *CPT1* forward primer (5′- GCTGTCTCATCATTTTGGCAAAGCGAATAATTCTAGGGTCGAC-3′) and *CPT1* reverse primer (5′- GTCTTTGTAGTCGGTACCAGG-3′), *rtTA* segment was generated using *CMV-Tet3G* (Clontech) as a template and *CPT2* forward primer (5′- CCTGGTACCGACTACAAAGACTCTAGACTGGACAAGAGCAAAG-3′) and *CPT2* reverse primer (5′- ggcagcctgcacctgaggagtgcatgtctggatccttacttag-3′). These *pCAG* fragment, *loxP-nG-STOP-loxP-mC-P2A* fragment and *rtTA* segment were gel extracted and assembled as described above.

#### *(viii)* *pMBP-loxP-nG-STOP-loxP-rtTA-P2A-mC*

*pCAG-loxP-nG-STOP-loxP-rtTA-P2A-mC* plasmid was digested by *ScaI-EcoRI* and *nG-STOP-loxP-rtTA-P2A-mC* segment was collected by gel extraction. *pMBP-loxP-nG-STOP-loxP-mC-P2A-DTR* was digested by *ScaI-EcoRI* followed by phenol-chloroform extraction and ethanol precipitation. The DNA solution containing *ScaI-EcoRI* digested fragments (2651 bp, 2528 bp and 4189 bp) was further digested by *SphI* to make gel extraction easier (2651 bp, 1751 bp, 777 bp and 4189 bp), and the *pMBP-loxP* fragment (2651 bp) was collected using gel extraction. *pMBP-loxP* fragment and *nG-STOP-loxP-rtTA-P2A-mC* fragment were ligated with *ScaI-EcoRI* site.

#### *(ix)* *pMBP-loxP-nG-STOP-loxP-mC-P2A-rtTa*

*pMBP-loxP* fragment (2651 bp) was obtained as described above. *nG-STOP-loxP-mC-P2A-rtTA* fragment was collected by *ScaI-EcoRI* digestion of *pCAG-loxP-nG-STOP-loxP-mC-P2A-rtTA* and gel extraction. *pMBP-loxP* fragment and *nG-STOP-loxP-mC-P2A-rtTA* fragment were ligated with *ScaI-EcoRI* site.

#### *(x)* *PB-CAG-loxP-nG-loxP-mC-P2A-rtTA* construct and *PB-CAG-loxP-nG-loxP-rtTA-P2A-mC*

Fragment containing *SpeI* site, piggyBAC terminal repeat sequences and *BglII* site was PCR amplified using *PBCAG-mRFP* (Addene #40996) (Chen and LoTurco, 2012) and forward primer (5′-agctggacatcacctcccacaacg-3′) and reverse primer (5′-tagcactagtctcgatatacagatcgataa-3′). The fragment was digested by *SpeI* and *BglII*. *pCAG-loxP-nG-STOP-loxP-rtTA-P2A-mC* construct or *pCAG-loxP-nG-STOP-loxP-mC-P2A-rtTA* construct were digested with *SpeI* and *BglII*, and generated *SpeI-CAG-loxP-nG-STOP-loxP-rtTa-P2A-mC-BglII* segment or *SpeI-CAG-loxP-nG-STOP-loxP-mC-P2A-rtTA-BglII* segment were generated, respectively. The segments and the SpeI-piggyBAC terminal repeat sequences-BglII fragment were ligated to generate *PB-CAG-loxP-nG-loxP-mC-P2A-rtTA* construct and *PB-CAG-loxP-nG-loxP-rtTA-P2A-mC* construct.

#### *(xi) pMBP(M321)-loxP-nG-STOP-lox2272-DTR-loxP-lox2272*

The M3 region of mouse MBP promoter was PCR amplified from mouse genomic using *M3* forward primer (5′-ATGATGGACGTCGTGGCAGATTTAGACTCCTTACC-3′) and *M3* reverse primer: (5′-CCGCTAGACGTCAGCCTGGTTCTGGAGTTGC-3′) according to Dib et al. (2011) with modulating linker region to add *AatII* restriction sites (Dib et al., 2011). M3 region was inserted into *AatII* site of *pMBP-loxP-nG-STOP-lox2272-DTR-loxP-lox2272* plasmid.

#### *(xii) pMBP(M321)-loxP-nG-STOP-lox2272-DTR-loxP-lox2272-pMBP(M321)-SnaBI*

Isothermal assembly (Gibson et al., 2009) was used to generate this construct (Gibson et al., 2009). *pMBP(M321)-loxP-nG-STOP-lox2272-DTR-loxP-lox2272* plasmid was linearized by SphI. *pMBP(M321)* sequence was PCR amplified with *M321* forward primer (5′-cttgcatgaagcggccgcaagcatgGCATGCGACGTCGTGGCAGATTTAGA-3′) and *M321* reverse primer (5′-CAGTGACCCGGAATCTGCAGGCATGGCATGCTACGTAACCCTAGAATTATTCGAGCT-3′). *pMBP(M321)-loxP-nG-STOP-lox2272-DTR-loxP-lox2272* fragment and *pMBP(M321)* fragment were assembled by Gibson assembly reaction.

#### *(xiii) pMBP(M321)-lox2272-nG-STOP-lox2272-mC-pMBP(M321)-SnaBI*

*pMBP(M321)-pMBP(M321)* fragment, *lox2272* fragment, *nG-STOP-lox2272* fragment and *mC-beta globlin poly A* fragment were assembled by isothermal assembly technique (Gibson et al., 2009) to generate this construct. *pMBP(M321)-loxP-nG-STOP-lox2272-DTR-loxP-lox2272-pMBP(M321)-SnaBI* was digested with *SalI* and 2762 bp fragment containing *loxP-nG-STOP-lox2272-DTR* and 8334 bp fragment containing *pMBP(M321)-pMBP(M321)* fragment were collected. *Lox2272* fragment was PCR amplified using the 2762 bp fragment containing *loxp-nG-STOP-lox2272-DTR* as a template and forward primer (5′ - AAGCTCGAATAATTCTAGGGAAGTAAGCTTGGGCTGCAGG-3′ ) and reverse primer (5′ - gctatgactacgaattgcgatTGAGGATCATCAAGCTTAGA-3′ ). The *nG-lox2272* fragment was PCR amplified using *pMBP(M321)-loxP-nG-STOP-lox2272-DTR-loxP-lox2272-pMBP(M321)-SnaBI* construct as a template and foward primer (5′ - atcgcaattcgtagtcatagcGCCACCATGCCAGAGCCAGC-3′ ) and reverse primer (5′ - ATCAAGCTTAGATCTCATATG-3′ ). The *mC-beta globlin poly A* fragment was PCR amplified using *pCAG-loxP-nG-STOP-loxP-rtTA-P2A-mC* plasmid as a template and forward primer (5′ - CATATGAGATCTAAGCTTGATGATATCGCCACCatgggttgctgtttctccaag-3′ ) and reverse primer (5′ - AGGATCGATCGGGATCCCGGTCTAGActcccatatgtccttccgag-3′ ).

***(xiv) pMBP(M321)-lox2272-nG-STOP-lox2272-mC-loxP-STOP-loxP-rtTA***

*pMBP(M321)-lox2272-nG-STOP-lox2272-mC* fragment was PCR amplified using *pMBP(M321)-lox2272-nG-STOP-lox2272-mC-pMBP(M321)-SnaBI* plasmid and forward primer (5′ - AtccagctcgaccaagcttgTCCACAGAATCAGGGGATA-3′ ) and reverse primer (5′ - tcggcgcgttcgtactgttc-3′ ). P2A fragment was PCR amplified using *pCAG-loxP-nG-STOP-loxP-mC-P2A-rtTA* as a template and forward primer (5′ - gaacagtacgaacgcgccga-3′ ) and reverse primer (5′ - ATTTCCtttgtagtcggtaccaggtc-3′ ). DNA fragment containing *loxP-3xpolyA-loxP* was PCR amplified using *Ai65(RCFL-tdT)* targeting vector (Madisen et al., 2015) (Addgene# 61577) as a template and forward primer (5′ - CTAGGGAAGAAGAGAGACCCAGGaaatataacttcgtataat-3′ ) and reverse primer (5′ - GTCCTTGTATTTCCGAAGACAgcaggtcgagggacctaata-3′ ). This *loxP-3xpolyA-loxP* sequence was PCR amplified using forward primer (5′ - gtaccgactacaaaGGAAATataacttcgtat-3′ ) and reverse primer (5′ - tccagtctagaAATGACgggacctaataacttcgtat-3′ ) to add DNA sequence compatible to the other segments, and named *loxP-STOP-loxP*. *rtTA-SV40polyA* fragment was PCR amplified using *pCMV-Tet3G* as a template and forward primer (5′ -CCCGTCATTtctagactggacaagagcaa-3′ ) and reverse primer (5′ - caagcttggtcgagctggat-3′ ). These *pMBP(M321)-lox2272-nG-STOP-lox2272-mC* fragment, *P2A* fragment, *loxP-STOP-loxP* fragment and *rtTA-SV40polyA* fragment were assembled using isothermal Gibson assembly reaction to generate *pMBP(M321)-lox2272-nG-STOP-lox2272-mC-loxP-STOP-loxP-rtTA* plasmid.
